# Structure of the human K_2P_13.1(THIK-1) channel reveals a novel hydrophilic pore restriction and lipid cofactor site

**DOI:** 10.1101/2024.06.26.600491

**Authors:** Shatabdi Roy-Chowdhury, Seil Jang, Fayal Abderemane-Ali, Fiona Naughton, Michael Grabe, Daniel L. Minor

## Abstract

The halothane-inhibited K_2P_ leak potassium channel K_2P_13.1 (THIK-1)^1–3^ is found in diverse cells^1,4^ including neurons^1,5^ and microglia^6–8^ where it affects surveillance^6^, synaptic pruning^7^, phagocytosis^7^, and inflammasome-mediated interleukin-1β release^6,8,9^. As with many K_2P_s^1,5,10–14^ and other voltage-gated ion channel (VGIC) superfamily members^3,15,16^, polyunsaturated fatty acid (PUFA) lipids modulate K_2P_13.1 (THIK-1)^1,5,14,17^ via a poorly understood mechanism. Here, we present cryo-electronmicroscopy (cryo-EM) structures of human K_2P_13.1 (THIK-1) and mutants in lipid nanodiscs and detergent. These reveal that, unlike other K_2P_s^13,18–24^, K_2P_13.1 (THIK-1) has a two-chamber aqueous inner cavity obstructed by a M4 transmembrane helix tyrosine (Tyr273, the flow restrictor). This hydrophilic barrier can be opened by an activatory mutation, S136P^25^, at natural break in the M2 transmembrane helix and by intrinsic channel dynamics. The structures also reveal a buried lipid in the P1/M4 intersubunit interface at a location, the PUFA site, that coincides with the TREK subfamily K_2P_ modulator pocket for small molecule agonists^18,26,27^. This overlap, together with the effects of mutation on K_2P_13.1 (THIK-1) PUFA responses, indicates that the PUFA site lipids are K_2P_13.1 (THIK-1) cofactors. Comparison with the PUFA-responsive VGIC Kv7.1 (KCNQ1)^28–31^ reveals a shared role for the equivalent pore domain intersubunit interface in lipid modulation, providing a framework for dissecting the effects of PUFAs on the VGIC superfamily. Our findings reveal the unique architecture underlying K_2P_13.1 (THIK-1) function, highlight the importance of the P1/M4 interface in control of K_2P_s by both natural and synthetic agents, and should aid development of THIK subfamily modulators for diseases such as neuroinflammation^6,32^ and autism^6^.

## Introduction

K_2P_ channels comprise six subfamilies that conduct leak potassium currents essential for membrane potential stabilization and cellular excitability regulation^3,16^ and influence diverse normal and pathophysiological processes^2,3^ including: action potential propagation^33,34^, pain^35–37^, sleep^38^, intraocular pressure^39^, retinal visual processing^40^, migraine^41^, depression^42^, pulmonary hypertension^43^, sleep apnea^44^, and inflammasome activation in macrophages^8,45,46^ and microglia^6,8,47^. Unlike other potassium channel classes, the selectivity filter (SF) ‘C-type’ gate forms the principal site of K_2P_ modulation^12,13,18,27,48–50^. Structural studies have shown that the TREK^18–20^, TWIK^21,22^, TALK^23^, and TASK^24^ subfamilies share a common architecture^13^. However, the THIK subfamily, comprising K_2P_13.1 (THIK-1) and K_2P_12.1 (THIK-2), has remained structurally uncharacterized. This subfamily is thought to have a role in anesthesia^51^, apoptosis^52^, and ischemic responses^17^, is inhibited by halothane^1,5,25^, lidocaine^5^, bupivacaine^5^, quinidine^5^, and the phosphodiesterase inhibitor 3-isobutyl-1-methyl-xanthine (IBMX)^53^, and is activated by the signaling lipid arachidonic acid^1,5^. Recently, K_2P_13.1 (THIK-1) has emerged as a key player in microglial function affecting ramification and surveillance^6^, synapse pruning^7^, and NLRP3 (<u>N</u>OD, <u>L</u>RR and <u>p</u>yrin domain-containing protein <u>3</u>) inflammasome-mediated release of interleukin 1β (IL-1β)^6,8,9^, making it a target for the development of agents against neuroinflammation^32^.

To understand THIK function, we determined the structure of human K_2P_13.1 (THIK-1) and mutants in lipid nanodisc and detergent environments using cryo-EM. The structures reveal a set of features unique to the THIK subfamily. A hydrophilic residue, Tyr273, the flow restrictor, forms a barrier that divides the central aqueous cavity into two parts: the cytoplasmic vestibule and a small aqueous chamber below the SF, termed the ‘pond’. Structures bearing the S136P^25^ activatory mutation show that this change opens the flow restrictor by exploiting the natural flexibility of a M2 transmembrane helical break present in all K_2P_s. The structures also reveal a polyunsaturated fatty acid (PUFA) bound in the P1/M4 helix intersubunit interface, the PUFA site, coordinated by an unusual, stacked arginine pair, the arginine sandwich, in which mutation sensitizes the channel to lipid modulation. Strikingly, the THIK PUFA site overlaps with the TREK subfamily K_2P_ modulator pocket where synthetic small molecule activators bind^13,18,26,27^. This structural and functional commonality establishes the role of the P1/M4 interface in natural and synthetic ligand control of K_2P_ function. Natural ligands such as fish and plant oil ω3 and ω6 PUFAs affect many voltage-gated ion channel (VGIC) superfamily members at binding sites that are thought to include ones located in the pore domain (PD)^15^. Comparison of the K_2P_13.1 (THIK-1) PUFA site with the proposed PD PUFA site on the cardiac I_Ks_ channel Kv7.1 (KCNQ1)^28–31^ uncovers structural commonalities that define a framework for understanding PUFA action in diverse VGIC superfamily members and highlight the potential of this shared PD intersubunit interface as a site for modulator development.

## Results

### Structure of human K_2P_13.1 (THIK-1) in lipid nanodiscs reveals novel features

We identified a human K_2P_13.1 (THIK-1) construct spanning residues 1-350 bearing N59Q and N65Q mutations to remove glycosylation, denoted K_2P_13.1_EM_, that maintained native-like function (Figs. S1a-c) and that could be readily purified using heterologous expression in HEK293 cells. Incorporation of K_2P_13.1_EM_ into brain polar lipid nanodiscs (Figs. S1d-e) for single particle cryo-electronmicroscopy (cryo-EM) (Figs.S1f-j) yielded a high-quality density map having an overall resolution of 2.65Å (Figs. S2a-b, Table S1) and local resolution as good as 2.4Å (Fig. 2c) that define the K_2P_13.1_EM_ structure (Figs. 1a and S2a-d). K_2P_13.1_EM_ shows the canonical K_2P_ structure (Figs. 1a and S2a)^13^, comprising a homodimer in which the M1 and M3 transmembrane helices from PD1 and PD2 face the bilayer, while M2 and M4 transmembrane helices from PD1 and PD2 line the central ion conducting pore. As in other K_2P_s^18–24^, the M1 helices are domain swapped and connect to an extracellular domain, the Cap, that sits above the extracellular mouth of the selectivity filter (SF)^13^. Consistent with the determination of the structure under high potassium conditions (200 mM KCl)^13,27^, potassium ions occupy the S1-S4 SF ion binding sites (Figs. 1a and S3a). Similar to exemplar structures of members of the TREK^18–20,27,34,54–60^, TWIK^21,22^, and TALK^23^ K_2P_ subfamilies, the intracellular face of the channel is open to the cytoplasm and lacks an intracellular gate (Figs. 1a-b and S3b-c).

**Figure 1.**
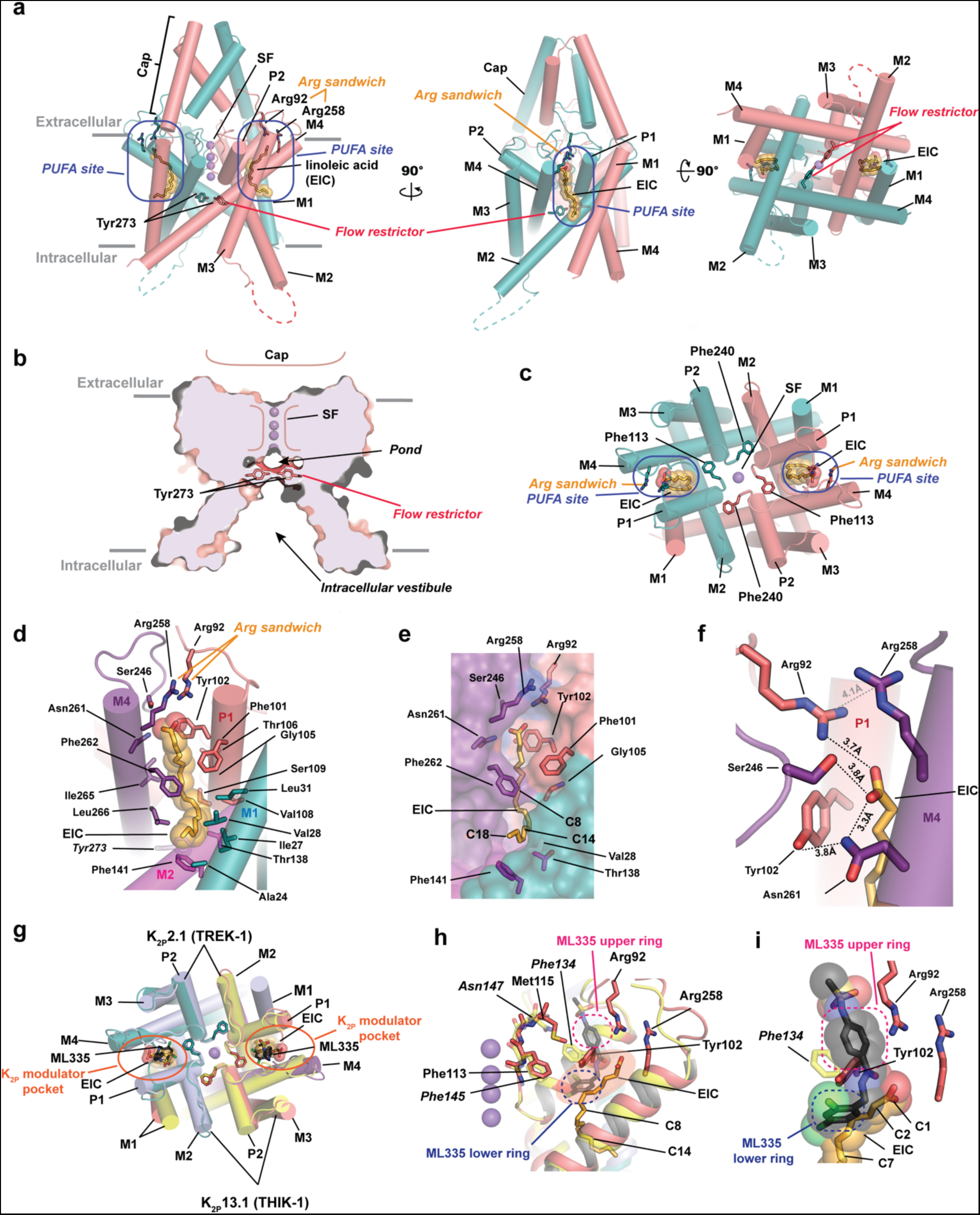
K_2P_13.1(THIK-1) structural features. **a,** K_2P_13.1(THIK-1) cartoon diagram (salmon and deep teal) showing side (left and center) and intracellular (right) views. Flow restrictor Tyr273 and Arg sandwich Arg92 and Arg258 residues are shown as sticks. PUFA site linoleic acid (EIC) is shown in space filling. Grey bars indicate membrane. Potassium ions are shown as purple spheres. **b,** Slice through the K_2P_13.1(THIK-1) structure showing the locations of the intracellular vestibule, flow restrictor, pond, and selectivity filter (SF). **c,** K_2P_13.1(THIK-1) extracellular view. M1, M2, M3, M4, P1, and P2 helices are labeled. SF residues Phe113 and Phe240 are shown as sticks. Other elements are displayed as in ‘a’. Cap is not shown. **d**, Details of the K_2P_13.1(THIK-1) PUFA binding site. M1 (teal), M2 (magenta), P1 (salmon), and M4 (violet) helices are shown as cylinders. PUFA site residues are shown as sticks and are labeled. Flow restrictor Tyr273 is in italics. EIC is shown as sticks and transparent space filling atoms. **e**, Surface view of ‘d’. Select residues are labeled. EIC C8, C14, and C18 carbons are indicated. Colors are as in ‘d’. **f**, K_2P_13.1(THIK-1) PUFA binding site hydrophilic interactions. Hydrogen bond network (black) surrounding the EIC acidic headgroup are shown. Arginine stack distance (grey) is indicated. **g,** and **h,** Structural superposition of K_2P_13.1(THIK-1) (salmon and deep teal) and the K_2P_2.1(TREK-1):ML335 complex (PDB:6CQ8) (yellow and light blue)^18^. **g,** Extracellular view showing position of the K_2P_2.1(TREK-1) K_2P_ modulator pocket (orange oval). ML335 is shown as black sticks. EIC is shown as sticks and transparent space filling atoms. SF phenyalanine sidechains are shown as sticks. M1, M2, M3, M4, P1, and P2 helices are labeled. **h,** Side view showing overlap regions of K_2P_ modulator pocket and PUFA site (orange). ML335 upper (pink) and lower (blue) rings are indicated. EIC C8 and C14 carbons are indicated. K_2P_2.1(TREK-1) residues are in italics. **i,** Closeup view of ‘h’

**Figure 2.**
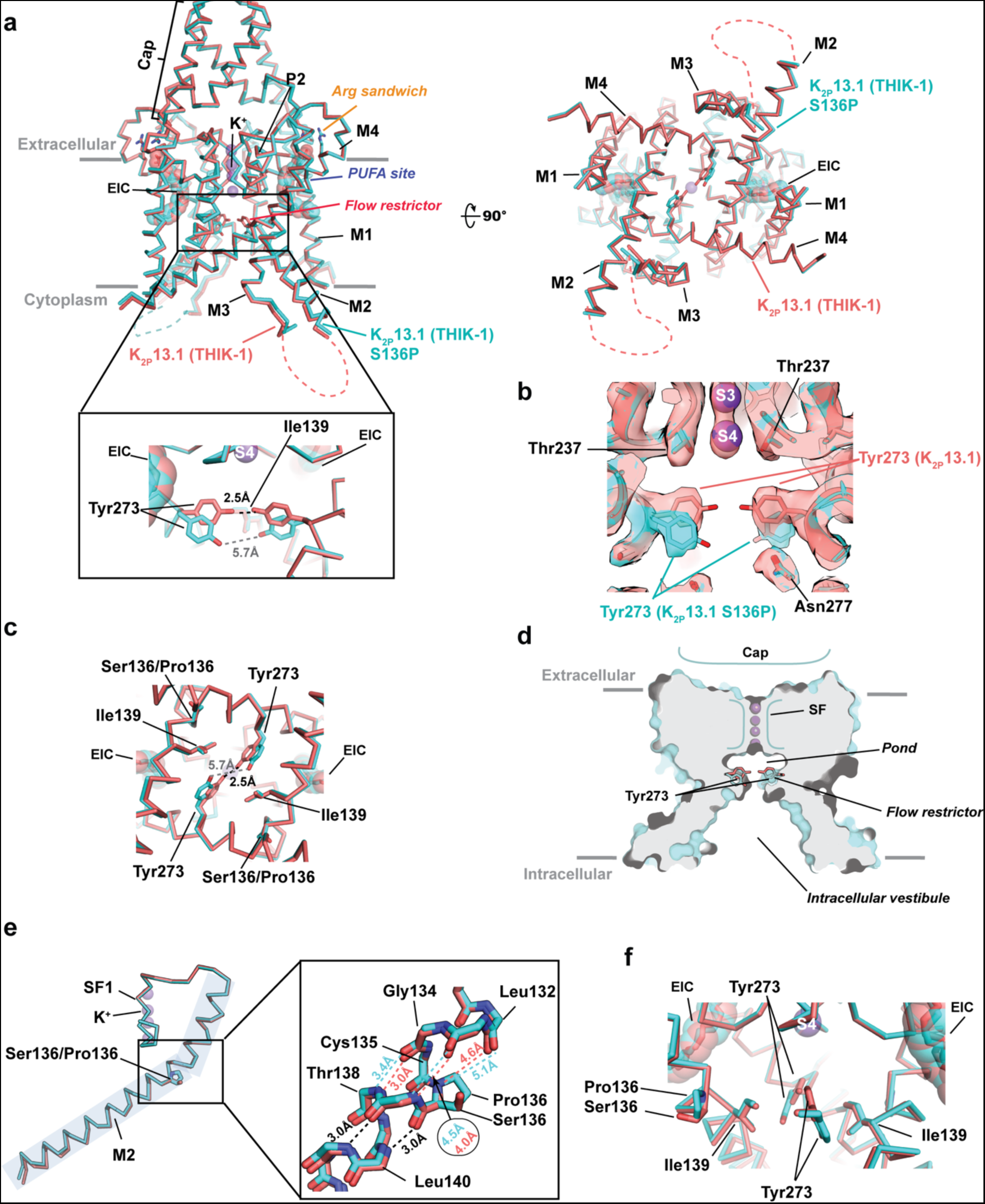
S136P mutation widens the K_2P_13.1 (THIK-1) flow restrictor. **a,** Ribbon diagram superposition of the structures of K_2P_13.1 (THIK-1) (salmon) and K_2P_13.1 (THIK-1) S136P (cyan). Flow restrictor Tyr273 and Arg sandwich Arg92 and Arg258 residues are shown as sticks. PUFA site linoleic acid (EIC) is shown in space filling. Inset shows closeup view of flow restrictor. Cytoplasmic view is shown on the right. Potassium ions are purple spheres. **b,** Cryo-EM density (8σ) for flow restrictor regions of K_2P_13.1 (THIK-1) (salmon) and K_2P_13.1 (THIK-1) S136P (cyan). **c,** Cytoplasmic view of flow restrictor conformational changes. **d,** Slice through the K_2P_13.1(THIK-1) S136P structure showing the locations of the intracellular vestibule, flow restrictor, pond, and selectivity filter (SF). K_2P_13.1 (THIK-1) Tyr273 conformation (salmon) is shown as sticks. **e,** Structural comparison of S136P mutation site. Inset shows local conformation and hydrogen bond distances for K_2P_13.1 (THIK-1) (salmon) and K_2P_13.1 (THIK-1) S136P (cyan). Common distances are in black. **f,** Zoomed in view of S136P site. Figure is ∼90° rotated relative to the view in the ‘a’ inset.

The K_2P_13.1 overall topology conforms with the general K_2P_ architecture but displays prominent variations from each of the other subfamilies (Fig. S4). K_2P_13.1 M4 is near the ‘up’ position seen in some TREK structures^18^ (Figs. S4a-b), but K_2P_13.1 M2 and M3 are closer to the central axis, making for a narrower channel profile when viewed from the cytoplasmic side (Figs. S4a-b). Similarly, K_2P_13.1 has a narrower cytoplasmic profile than the TALK representative K_2P_5.1 (TASK-2)^23^ due to differences in the positions and helical pitches of M1, M3 and M4 (Fig. S4c). By contrast, the pitch and positions of all four K_2P_13.1 transmembrane helices also diverge from those in K_2P_1.1 (TWIK-1)^21^ (Fig. S4d) making for a wider inner pore in K_2P_13.1 (THIK-1). K_2P_13.1 (THIK-1) lacks the cytoplasmic X-gate seen in K_2P_3.1 (TASK-1)^24^ and also diverges in the positions of its M3 and M4 (Fig. S4e). Hence, the K_2P_13.1_EM_ structure highlights the structural diversity within the general K_2P_ architectural framework.

In addition to these overall K_2P_ architecture variations, the K_2P_13.1_EM_ structure revealed two unexpected features conserved throughout K_2P_13.1(THIK-1) homologs from vertebrates spanning from amphibians to humans (Fig. S5). First, the K_2P_13.1 inner pore cavity is divided into two chambers by a barrier formed by the Ile139 and Tyr273 sidechains in which the Tyr273 hydroxyl groups make a hydrogen bond across the narrowest part (2.5Å) (Figs. 1a-b and S3a, S6a-b). These tyrosines form the deepest part of this barrier with respect to the cytoplasmic vestibule (Figs. S6b and S7a) and are unique to the THIK subfamily (Fig. S6c). We denote Tyr273 as the ‘flow restrictor’ as unlike classical hydrophobic gates in many VGIC superfamily members^61^, the Tyr273 pair forms a barrier comprising hydroxyl moieties that have potential to coordinate passing ions. The flow restrictor is much closer to the SF than the inner gates in other channels (∼7Å vs. ∼20Å from the S4 ion to the K_2P_13.1 flow restrictor and K_2P_3.1 cytoplasmic X-gate^24^, respectively) and divides the central aqueous pore into a small chamber between the flow restrictor and SF (denoted ‘the pond’) and large intracellular vestibule bounded by an asparagine ring (Asn143 and Asn277) found in both THIK subfamily members (Figs. S6a-d). The pond has a volume of ∼20 water molecules and maximal dimensions of ∼4.5Å along the channel central axis x 20Å across and is electrostatically negative (Fig. S7b) in contrast to the electrostatically positive cytoplasmic vestibule. This two-chambered central cavity sets the THIK subfamily apart from other K_2P_s.

The second K_2P_13.1 (THIK-1) unexpected feature is a well-defined banana-shaped density in the interdomain interface formed by the P1 and M4 helices of each monomer (Figs. S3b-c). Based on its shape, we modeled this density as linoleic acid (EIC) (Figs. S2b and S3c), a polyunsaturated fatty acid (PUFA) from a class of lipids that affects many K_2P_s^1,5,10–14^ and thus, denoted this site as the ‘PUFA site’ (Fig. 1a). Due to steric considerations, the PUFA site lipid requires a largely straight chain between the carboxylate and C9 position and an unsaturated bond at C12 to accommodate the bend in the density. Accordingly, the density could be equally well fit by α-linolenic acid, which has an additional unsaturated bond at C15. Determination of the structure of K_2P_13.1_EM_ in detergent micelles at 2.95Å resolution (Figs. S8-S9, Table S1), revealed a structure that is not different from the K_2P_13.1_EM_ nanodisc structure (RMSD = 0.248Å) except for having one more helical turn at the end of M2 (Arg163-Arg167) (Figs. S10a-c) and a weaker density in the SF for the S2 ion (Fig. S10d). Importantly, the same banana-shaped density was present in the PUFA site (Fig. S9a), indicating that this lipid originated from the host cell and not from the lipids used for nanodisc reconstitution. Both structures also show numerous common positions of hydrophobic chains in the outer membrane leaflet lipid belt that surrounds the channel (Figs. S2a and S2c, S9b, and S10e-f) and inner leaflet site between the M1 and M2 helices (Fig. S10e), suggesting that the outer surfaces of the channel strongly associate with lipids.

The PUFA is deeply embedded within the protein structure and contacts 18 amino acids from the M1, P1, M2, and M4 helices (Figs. 1c-e and S6c-g) of which the majority are hydrophobic. Residues from the SF2-M4 loop (Ser246), M4 helix (Asn261), and loop connecting the Cap and P1 helix (Arg92) coordinate the PUFA headgroup via a set of hydrogen bonds (Ser246 and Asn261) and a salt bridge (Arg92) (Figs. 1f and S6e). Arg92 is part of an unusual structure in which its sidechain guanidinium group forms a 4.0Å face-to-face stack (denoted ‘the arginine sandwich’) with the guanidinium of Arg258 from M4 (Fig. 1a, d, and f). Such arginine-arginine stacks have been observed in a wide variety of proteins where they are thought to affect molecular recognition, protein-protein, and protein-ligand interactions^62,63^. This arginine sandwich pair occurs only in the THIK subfamily (Figs. S6c and f), making it, together with the flow restrictor tyrosine, signature features of this K_2P_ subfamily.

The P1/M4 interface is central to function of the K_2P_ SF gate^13,18,27,48,56,64^ and houses the K_2P_ modulator pocket, a small molecule binding site in the TREK subfamily where the synthetic activators ML335 and ML402 bind^13,18,26,27^. Superposition of K_2P_13.1 (THIK-1) and the K_2P_2.1(TREK-1):ML335 complex^18^ (Figs. 1g-h) shows that the K_2P_13.1 PUFA site and K_2P_ modulator pocket coincide. The positions of the PUFA carboxylate and C2-C7 carbons partially overlap with the portion of the K_2P_ modulator pocket occupied by the ML335 activator inter-ring linker and lower ring (Figs. 1h-i). A subset of these PUFA elements interact with the K_2P_13.1 P1 helix residue Tyr102, a residue that corresponds to K_2P_2.1(TREK-1) Phe134 that provides a buttress position for K_2P_ modulator pocket small molecule activator binding^18,26^ (Fig. 1h-i). Arg92 from the K_2P_13.1 arginine sandwich occupies the upper part of the K_2P_ modulator pocket near the position corresponding to the ML335 upper ring (Fig. 1h-i). The other arginine sandwich member, K_2P_13.1 Arg258, is stacked on top of Arg92, and corresponds to K_2P_2.1(TREK-1) Lys271 that makes cation-π interactions with the modulator upper ring^18,26^. Hence, even though the ligands P1/M4 interface ligands from K_2P_13.1(THIK-1) and K_2P_2.1(TREK-1) have different physicochemical properties, the two binding sites bear similarities that support the idea that the THIK subfamily PUFA site is a variation of the TREK subfamily K_2P_ modulator pocket.

One key difference between the PUFA and K_2P_ modulator sites is the interactions made by the lower part of the PUFA for which the TREK modulators have no corresponding element. The K_2P_13.1(THIK-1) PUFA C8 to C14 methylene groups snake along the P1/M4 interface in a manner that allows the C15-C18 carbons to emerge through a gap between the K_2P_13.1 P1, M2, and M4 helices below residues Phe101, Gly105, and Ile265 at the middle level of the lipid bilayer (Figs. 1d-e). K_2P_2.1(TREK-1) lacks this gap due to the presence of a large amino acid, Phe170, at the K_2P_13.1 Thr138 position that forms the lower bound of the opening (Figs. 1d-e and S6e). Together, these observations identify the K_2P_13.1 (THIK-1) PUFA site as a variation of the TREK subfamily K_2P_ modulator pocket, reveal how this interface pocket is tuned to accommodate diverse ligands in different K_2P_ subfamilies, and show that both natural and synthetic ligands can target this site.

### An activatory mutation opens the K_2P_13.1 (THIK-1) flow restrictor

All K_2P_s except for the THIK subfamily have a proline at a position where a break occurs in the M2 helical structure^21,25,58^(Fig. S6d). Placing a proline at this position in K_2P_13.1 (THIK-1) and K_2P_12.1 (THIK-2), S136P and A155P, respectively, increases THIK channel currents in *Xenopus* oocytes and mammalian cells^25^. To understand how this mutation acts, we made the S136P mutation in the K_2P_13.1_EM_ background, validated that this change increases channel function as expected^25^ (Figs. S11a-c), and used cryo-EM to determine its structure in lipid nanodiscs (2.36Å) (Figs. S11d-j and S12, Table S1) and detergent environments (2.73Å) (Figs. S13-S14, Table S1). Both are similar (all atom RMSD = 0.486Å) apart from an extra five residue extension of the M2 helix (residues 163-167) (Figs. S15a-c) and weaker density for the SF S2 ion (Figs. S15d-e) in the detergent environment, recapitulating the K_2P_13.1_EM_ nanodisc and detergent structural differences (Figs. S3a and S10a-b and d). Overall, K_2P_13.1_EM_ S136P is also very similar to K_2P_13.1_EM_ (RMSD_Cα_ = 0.352Å) (Fig. 2a). There are ions at SF sites S1-S4 and density matching EIC coordinated by arginine sandwich in both PUFA binding sites (Fig. 2a). However, inspection of the flow restrictor revealed that Tyr273 tips away from the channel central axis increasing the phenolic oxygen interatom distance from 2.5Å to 5.7 Å (Fig. 2b-c). This local conformational change opens a gap providing an aqueous pathway between the pond and inner vestibule (Fig. 2d, Movies S1 and S2) and suggests that Tyr273 movement is central to the mechanism by which the S136P mutation increases channel function.

Remarkably, despite the sidechain conformational changes at Tyr273, the backbones and Cβ atoms of Ser136 and S136P from the K_2P_13.1_EM_ and K_2P_13.1_EM_ S136P structures overlay extremely well (Figs. 2e-f). There is a small expansion of M2 kink due to changes in the Leu132-Gly134 backbone carbonyl positions (∼0.5Å) (Fig. 2e, Movies S1 and S2). Superposition of K_2P_13.1 (THIK-1) with TREK, TWIK, TASK, and TALK subfamily representatives where the corresponding Ser136 site is naturally a proline (Fig. S6d) shows that the SF1, SF1-M2 linker, and upper part of S2 before the kink align. By contrast, the lower part of M2 following the Ser136 site adopts a range of positions that span a ∼15° arc (Fig. S6h), indicating that this section of M2 is mobile. K_2P_13.1_EM_ and K_2P_13.1_EM_ S136P occupy the middle of this distribution. The K_2P_13.1 (THIK-1) Ser136 site occurs at a break at the N-terminus of a helical segment in all K_2P_ structures. Such sites accommodate prolines^65^. As a result, subtle changes in M2 caused by the S136P mutation appear to exploit the flexibility at this junction and allow M2 to flex in a way that enables the flow restrictor to open (Movies S1 and S2) and enhance function^25^.

### Flow restrictor shapes the energetic landscape of the permeant ion

The K_2P_ 13.1 (THIK-1) ion conduction pathway has a very different shape than the archetypal K_2P_ channel K_2P_2.1 (TREK-1) (Figs. 3a-c). Whereas the pore cavity below the K_2P_2.1 (TREK-1) SF is wide (Fig. 3c), the K_2P_ 13.1 (THIK-1) flow restrictor and Asn ring respectively block and narrow the central pathway below the SF (Fig. 3a). Although this general cavity shape is maintained the S136P mutant where the flow restrictor Tyr273 has been opened, the movement of Tyr273 not only connects but reshapes the sizes of the pond and vestibule (Figs. 3b and d). To test the functional importance of the flow restrictor constriction, we made a mutant, Y273F, that removed the Tyr273 hydroxyl that blocks the pathway (Figs. 1a-b and 2a). This change increased K_2P_ 13.1 (THIK-1) activity to a level similar to the S136P change (I/I_WT_=2.16±0.15 and 2.36±0.17, for Y273F and S136P, respectively) (Figs. 3e-f, Table 1). Reducing the flow restrictor size further with Y273L and Y273A yielded even larger current enhancements (I/I_WT_=4.17±0.20 and 4.85±0.23, respectively) (Figs. 3e-f, Table 1), supporting the idea that the Tyr273 flow restrictor is a key controller of K_2P_ 13.1 (THIK-1) activity. The M2 pore-lining residue Ile139 supports Tyr273 (Figs. S6a-b). Changing this position to alanine enhanced currents similar to Y273A (I/I_WT_=4.77±0.25 and 4.85±0.23, respectively) (Figs. 3e-f) consistent with the increased function reported for a glycine change at this site^14^. Rather than producing additive effects, combining I139A and Y273A was less stimulatory than the individual mutants (I/I_WT_=3.78 ±0.10) (Figs. 3e-f, Table 1), consistent with the structural interactions between these two residues. Similar trends were observed in the S136P background (Table 1). Y273F/L/A and I139A all increased currents more than S136P alone (I/I_S136P_=2.45±0.16, 3.25±0.26, 3.30±0.28 and 3.00±0.26, for Y273F, Y273L, Y273A, and I139A, respectively) (Figs. S16a-b, Table 1) and there was no additive effect of I139A/Y273A (I/I_S136P_=3.12±0.16) (Figs. S16a-b, Table 1). Thus, regardless of context, reducing the size of the flow restrictor residue or perturbing its nearest neighbor, dramatically increased channel function. Together, these data indicate that the flow restrictor position Tyr273 limits ion flux through the channel.

**Figure 3.**
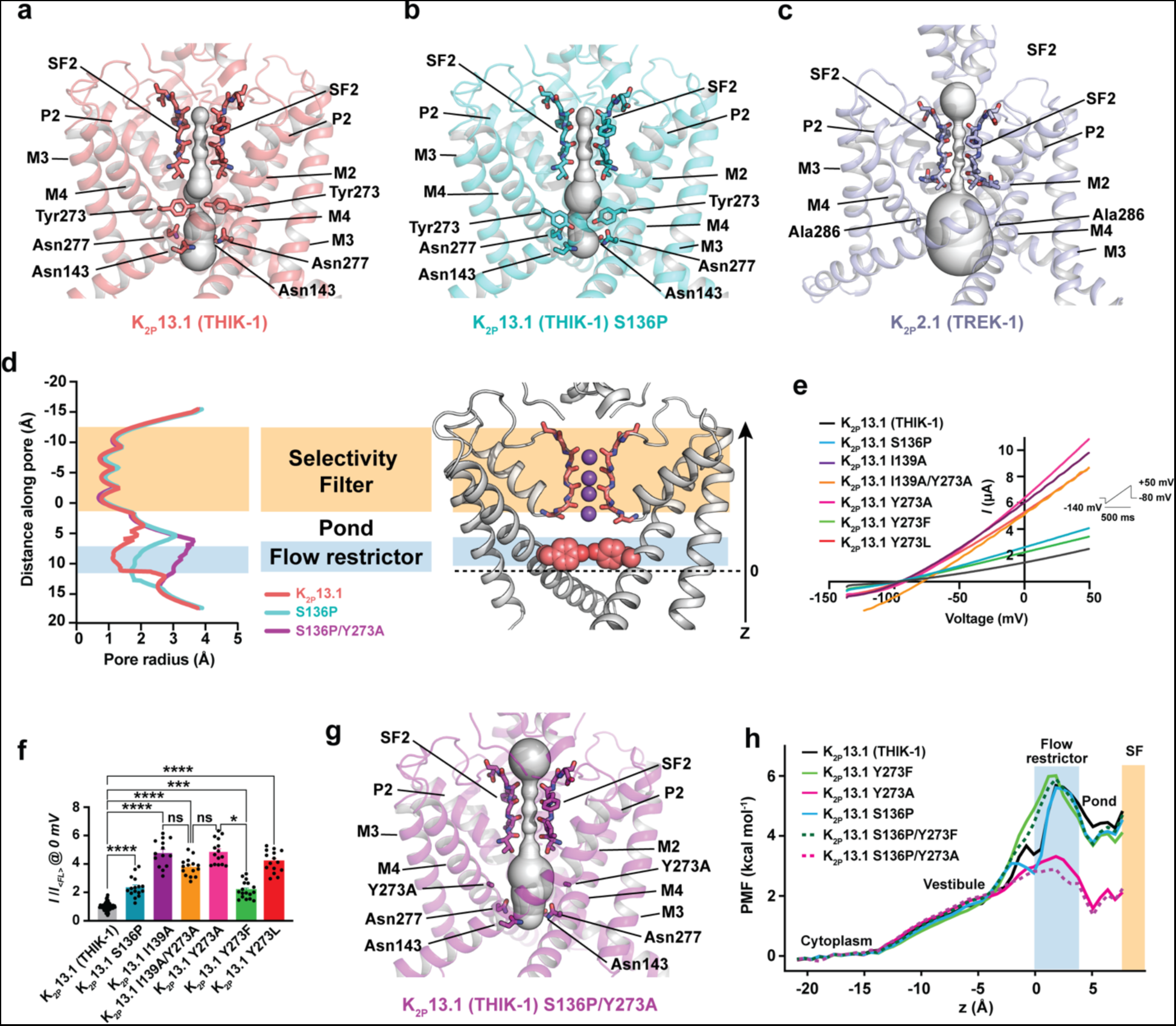
K_2P_13.1 (THIK-1) flow restrictor controls inner cavity shape and function. **a-c,** Pore profile comparisons for **a,** K_2P_13.1 (THIK-1) (deep salmon) **b,** K_2P_13.1 (THIK-1) S136P (cyan), and **c,** K_2P_2.1 (TREK-1) (PDB:6CQ6) (light blue)^18^ calculated using HOLE^102^. Selectivity filter 2 (SF2), Flow restrictor, and Asn ring residues are shown as sticks. Select channel elements are labeled. **d,** K_2P_13.1 (THIK-1) model and pore profiles. **e,** Representative TEVC current-voltage responses for K_2P_13.1 (THIK-1) (black) and K_2P_13.1 (THIK-1) mutants: S136P (cyan), I139A (purple), I136A/Y273A (orange), Y273A (magenta), Y273F (green), and Y273L (red). Inset shows protocol. **f,** Average currents I/I_<FL>_ at 0 mV for K_2P_13.1 (THIK-1) (black) and K_2P_13.1 (THIK-1) mutants: S136P (cyan), I139A (purple), I136A/Y273A (orange), Y273A (magenta), Y273F (green), and Y273L (red) where I_<FL>_ = average currents for full length K_2P_13.1 (THIK-1). **** p <0.0001, *** p <0.001, ** p <0.01, * p = 0.01 −0.05, n.s. p >0.05. **g,** Pore profile of K_2P_13.1 (THIK-1) S136P/Y273A (magenta). **h,** Potential of Mean Force (PMF) for K_2P_13.1 (THIK-1) (black), K_2P_13.1 (THIK-1) Y273F (green), K_2P_13.1 (THIK-1) Y273A (magenta), K_2P_13.1 (THIK-1) S136P (cyan), K_2P_13.1 (THIK-1) S136P/Y273F (dashed green), and K_2P_13.1 (THIK-1) S136P/Y273A (dashed dark magenta). Z axis corresponds to that shown in ‘d’. Statistical analysis was performed using the Kruskal-Wallis test (nonparametric ANOVA) followed by Dunn’s multiple comparisons test. In ‘d’ and ‘h’ locations of the Flow restrictor (light blue) and Selectivity filter (SF) (orange) are indicated.

**Table 1:** Relative current amplitudes at 0 mV.

| **K<sub>2</sub>P13.1 (THIK-1)** |  |  |  | **K<sub>2</sub>P13.1<sub>EM</sub>** |  |  |
| --- | --- | --- | --- | --- | --- | --- |
|  |  | *I*/*I*<sub>WT</sub> | **n** | *I*/*I*<sub>WT</sub> | *I*/*I*<sub>S136P</sub> | **n** |
|  | WT | 1.00 ± 0.02 | 107 | 0.93 ± 0.13 |  | 15 |
|  | S136P | 2.36 ± 0.17 | 16 | 1.39 ± 0.09 | 1.00 ± 0.06 | 28 |
| **Arg sandwich** | R92A | 0.74 ± 0.06 | 27 |  |  |  |
|  | R258A | 1.18 ± 0.07 | 30 |  |  |  |
| **Flow restrictor** | I139A | 4.77 ± 0.25 | 14 | 4.35 ± 0.38* | 3.00 ± 0.26 | 3 |
|  | I139A/Y273A | 3.78 ± 0.16 | 15 | 4.52 ± 0.37* | 3.12 ± 0.26 | 3 |
|  | Y273A | 4.85 ± 0.23 | 15 | 4.85 ± 0.43* | 3.30 ± 0.28 | 14 |
|  | Y273F | 2.16 ± 0.15 | 17 | 3.55 ± 0.23* | 2.45 ± 0.16 | 13 |
|  | Y273L | 4.17 ± 0.31 | 15 | 4.72 ± 0.38* | 3.25 ± 0.26 | 10 |
| **PUFA site** | F101A | 1.40 ± 0.11 | 25 |  |  |  |
|  | Y102A | 0.97 ± 0.08 | 25 |  |  |  |
|  | G105F | 0.57 ± 0.04 | 11 |  |  |  |
|  | G105I | 1.54 ± 0.24 | 9 |  |  |  |
|  | S246A | 1.32 ± 0.10 | 18 |  |  |  |
|  | N261A | 0.11 ± 0.01 | 14 |  |  |  |
|  | F262A | 1.39 ± 0.11 | 16 |  |  |  |
|  | R92A/N261A | 0.05 ± 0.01 | 6 |  |  |  |
|  | F101A/F262A | 1.52 ± 0.12 | 6 |  |  |  |

\* S136P background

To understand the consequences of flow restrictor disruption further, we determined the lipid nanodisc embedded structure of a mutant in which the flow restrictor residue was shorted to alanine, K_2P_13.1_EM_ S136P/Y273A, at 2.39 Å resolution (Figs. S17-S19). This structure is essentially identical to K_2P_ 13.1_EM_ S136P:ND (RMSD = 0.378Å) (Figs. S19a-d) and preserves the arginine sandwich, PUFA site lipid features (Fig. S19a) and four SF ions (Fig. S19e) of K_2P_13.1_EM_ and K_2P_13.1_EM_ S136P. The absence of the flow restrictor tyrosine (Figs. S19c-d) creates a continuous inner cavity (Fig. 3g) having a gap between Y273A β-carbons of 13.8Å (Fig. S19d) that is much larger than the 2.3Å and 5.7Å gaps between Tyr273 hydroxyls in the K_2P_ 13.1_EM_ (Fig. 2a) and K_2P_ 13.1_EM_ S136P (Figs. 2a and S19d) structures. As a result, the narrowest point is Ile139 (9.6Å) (Figs. 3d and S19c). This unobstructed path from the cytoplasm to SF together with the functional data for Y273A mutants (Figs. S16a-b and Table 1) further supports the idea that the flow restrictor and its movements control K_2P_13.1 (THIK-1) function.

To examine how the flow restrictor affects energetic barriers traversed by permeant ions, we computed the potentials of mean force (PMF) for potassium ion entry from the cytoplasm to the pond for K_2P_ 13.1 (THIK-1) and Y273F, Y273A, S136P, S136P/Y273F and S136P/Y273A mutants. These calculations show that ions entering from the cytoplasm must surmount a ∼6 kcal mol^-1^ barrier at the flow restrictor and that following entry into the pond are stabilized by a ∼2 kcal mol^-1^ exit barrier (Fig. 3h). The barrier height is similar for K_2P_13.1 (THIK-1), S136P, Y273F, and S136P/Y273F due to the fact that these simulations rapidly relax to the Tyr273 open conformation seen in the K_2P_13.1 (THIK-1) S136P (Fig. S20, Movies S3 and S4). This rapid relaxation supports the idea that the S136P mutation biases the Tyr273 conformation towards a state that is naturally sampled by the channel. Y273F also eliminates a minimum before the barrier that occurs due to coordination of potassium by Tyr273 hydroxyl groups (Fig. 3h), in line with the idea that this residue directly interacts with the permeant ion. Further, consistent with the wide intracellular opening in K_2P_13.1_EM_ S136P/Y273A (Figs. 3g, S19c-d), Y273A reduces the barrier between the cytoplasm and pond by ∼4 kcal mol^-1^ (Fig. 3h), explaining the stimulatory effects of Y273A versus Y273F (Fig. 3f). Taken together, these data support the idea that Tyr273 forms a barrier that limits ion passage from the cytoplasmic vestibule into the pond and can coordinate the permeant ion. Perturbation of this barrier affects channel function.

### PUFA site lipid is important for channel function

All K_2P_ 13.1 (THIK-1) structures have a lipid in the PUFA binding site (Figs. 1d, 2a, S10a, S15a, and S19a) that overlaps with point of ligand control in the TREK family (Figs. 1g-i)^13,18,26,27^. These structural observations, together with prior reports that the PUFA arachidonic acid could stimulate rat K_2P_ 13.1 (THIK-1)^1,5^ with a stoichiometry indicating two binding sites^1^ prompted us to investigate the roles of PUFAs in K_2P_ 13.1 (THIK-1) modulation by first testing the functional consequences of PUFA binding site mutations. Alanine substitution the arginine sandwich residue that directly coordinates the PUFA carboxylate, R92A (Fig. 1f), reduced channel function (I/I_WT_=0.74 ±0.06) (Figs. 4a-b, Table 1). Change of its partner, R258A (Fig. 1f) was mildly stimulating (I/I_WT_=1.18 ±0.07) (Figs. 4a-b, Table 1). Mutations at the other two PUFA carboxylate coordination sites, S246A and N261A (Fig. 1f), identified a particular importance for Asn261 as its removal eliminated channel function (I/I_WT_=1.32 ±0.10, 0.11 ±0.01, 0.05 ±0.01 for S246A, N261A, and R92A/N261A, respectively) (Figs. 4a-b, Table 1). Mutation of the buttress residue from the P1 helix, Y102A, was neutral (I/I_WT_=0.97 ± 0.08) (Table 1), whereas changes to residues framing the lateral membrane-facing portion of the PUFA site either enhanced (F101A, F262A, G105I, F101A/F262A) or decreased (G105F) function (Figs. 4a-b, Table 1). Placed in structural context, the data show that PUFA interactions with the residues that directly coordinate the lipid carboxylate (Arg92 and Asn261) are key for function, whereas the residues above (Ser246 and Arg258) or at the lateral opening to the membrane (Phe101, Gly105, Phe262) tune activity (Fig. 4c). Taken together, these data support the idea that the PUFA site is critical for K_2P_ 13.1 (THIK-1) activity.

**Figure 4.**
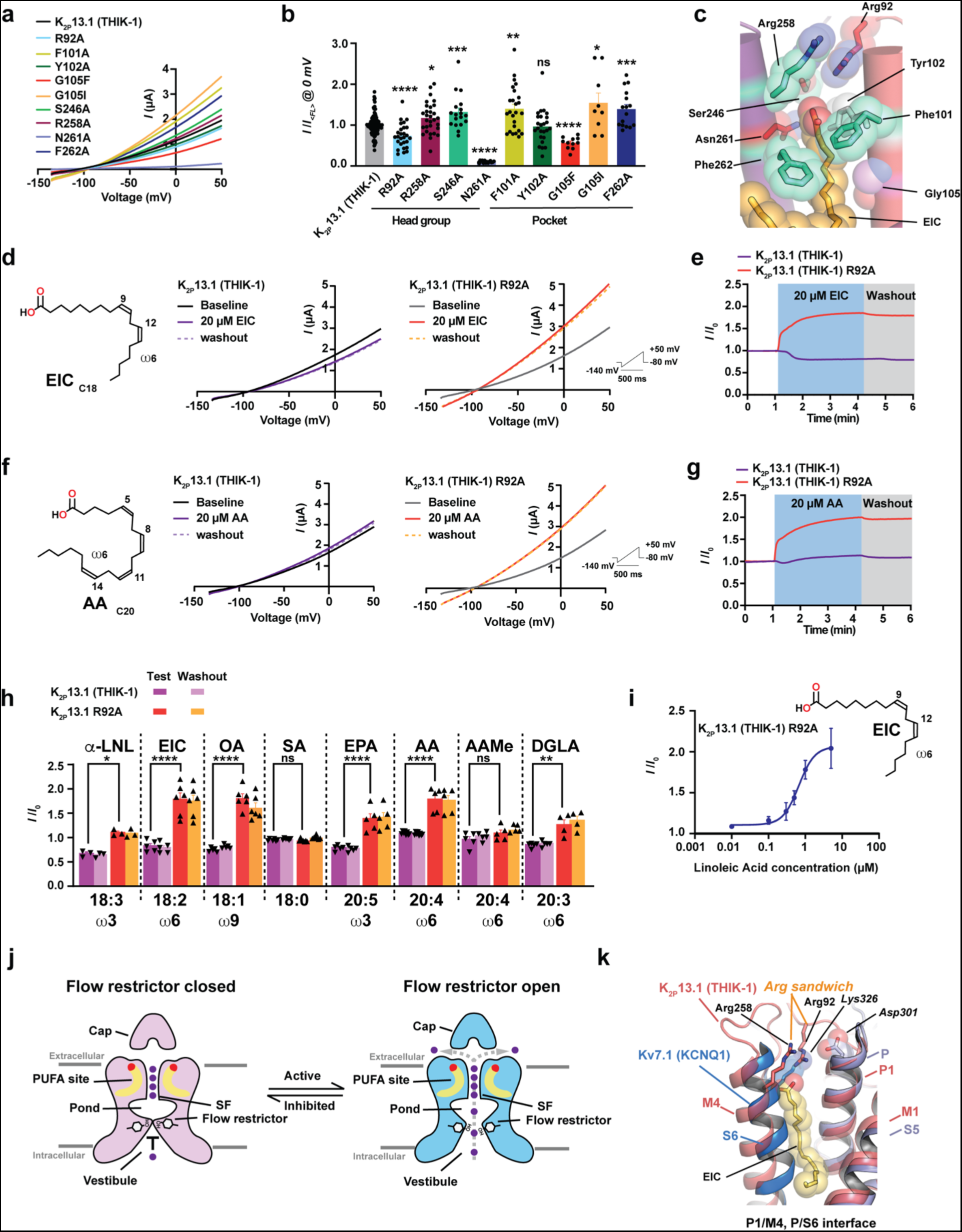
K_2P_13.1 (THIK-1) PUFA site is important for function. **a,** Representative TEVC current-voltage responses for K_2P_13.1 (THIK-1) (black) and K_2P_13.1 (THIK-1) mutants R92A (cyan), F101A (yellowgreen), Y102A (dark green), G105F (red), G105I (orange), S246A (green), R258A (maroon), N261A (blue), F262A (dark blue). Inset shows protocol. **b,** Average currents I/I_<FL>_ at 0 mV for K_2P_13.1 (THIK-1) (black) and K_2P_13.1 (THIK-1) mutants where I_<FL>_ = average currents for full length K_2P_13.1 (THIK-1). Colors are as in ‘a’. Lipid head group coordinating and pocket lining residues are indicated. **** p <0.0001, *** p <0.001, ** p < 0.01, * p = 0.01 −0.05. Statistical analysis was performed using the Mann-Whitney test. **c,** Effects of mutants on K_2P_13.1 (THIK-1) basal activity. Colors indicate increase (cyan), decrease (red), both (purple), or no effect. **d-g,** Representative TEVC current-voltage responses (**d** and **f**) and time course responses (**e** and **g**) for K_2P_13.1 (THIK-1) (purple) and K_2P_13.1 (THIK-1) R92A (red) to **d-e,** linolenic acid (EIC) and **f-g,** arachidonic acid (AA). **d,** and **f,** show lipid structure. Unsaturated bond locations are marked by numbers. Lipid class (ω6) is indicated. IV curves show baseline (black), lipid application (purple or red), and after 2’ washout (purple or orange dashed lines). **h,** Lipid modulation of K_2P_13.1 (THIK-1) and K_2P_13.1 (THIK-1) R92A I/I_0_ at 0 mV where I_0_ is the current before lipid application. Light purple and orange show response after 2’ washout. Lipids are: arachidonic acid (AA), arachidonic acid methyl ester (AAMe), eicosapentanoic acid (EPA), dihomo-γ-linolenic acid (DGLA), α-linolenic acid (α-LNL), linoleic acid (EIC), oleic acid (OA), and stearic acid (SA). X:Y indicates number of carbons and unsaturated bonds. **** p <0.0001, *** p <0.001, ** p < 0.01, * p = 0.01 −0.05, n.s. p >0.05. Statistical analysis was performed using the Kruskal-Wallis test (nonparametric ANOVA) followed by Dunn’s multiple comparisons test. **i,** TEVC dose response curve for K_2P_13.1 (THIK-1) R92A activation by linoleic acid. Curve shows fit to the Hill Equation. EC_50_= 0.7 µM, Hill coefficient ‘n’ = 2.0. **j,** Model of THIK function. Changes in the conformation equilibrium affect ion passage. **k,** Comparison of K_2P_13.1 (THIK-1) (deep salmon) and Kv7.1 (KCNQ1) (marine and light blue) (PDB:6UZZ)^103^ P1/M4 and P/S6 interfaces. Residues framing the putative Kv7.1 (KCNQ1) PUFA site^29^, Lys326 and Asp301 and EIC (yellow orange) are shown in space filling. Arginine sandwich pair is indicated.

As the PUFA site lipid is deeply buried in the P1/M4 interface and must be tightly bound to survive the purification required for structure determination, we decided to compare the effects of PUFA application to wild-type channels and the R92A mutant, as this change reduced basal activity (Figs. 4a-b, Table 1) and is expected to affect lipid binding. In line with the idea that the PUFA is strongly bound to the wild-type channel, we observed modest responses for K_2P_13.1 (THIK-1) to the C18 lipid that best fits the cryo-EM density (linoleic acid, EIC) (Figs. 4d-e), and arachidonic acid (AA) (Figs. 4f-g), a C20 lipid reported to simulate rat K_2P_ 13.1 (THIK-1)^1,5^. By contrast, these two ω6 PUFAs produced a robust and lasting activation of R92A (Figs. 4d-h). Further, arachidonic acid methylester (AAMe), the neutral version of AA, was ineffective (Figs. 4h and S21a-b), underscoring the importance of the negatively charged carboxylate. α-linolenic acid (α-LNL), eicosapentanoic acid (EPA), dihomo-γ-linolenic acid (DGLA), and oleic acid (OA) also activated R92A contrasting their minimal or slightly inhibitory effects on K_2P_13.1 (THIK-1) (Figs. 4h and 21c-j). These results demonstrate that unsaturated fatty acids of different lengths from the ω3 (α-LNL and EPA), ω6 (EIC, AA, DLGA), and ω9 (OA) classes can activate the channel. Importantly, stearic acid (SA), a C18 saturated fatty acid, had no effect (Fig. 4h, S21k-l). Measurement of the dose-response curve for K_2P_13.1 (THIK-1) R92A to linoleic acid revealed a saturating response (EC_50_ = 0.7µM) having a stoichiometry that matches the number of observed PUFA sites (Figs. 1a and 4i). Together with the mutational data, these R92A gain-of-function effects on lipid modulation suggest that the PUFA is largely a structural element rather than a regulatory lipid in the wild-type channel and that mutations that affect the PUFA site can sensitize the channel to lipid modulation.

To understand how the PUFA site might accommodate a range of PUFAs, we used molecular dynamics simulations in which we placed different PUFAs at the EIC position. In all cases, the helices forming the PUFA site accommodated the different lipids with minimal movement (Fig. S22a). By contrast, we saw varied behavior for the lipids. The C18 ω6 PUFA that fit the density best, EIC, was very stable, as was its C18 ω3 counterpart α-LNL (Fig. S22b). DGLA and AA, two C20 ω6 PUFAs, rose through the binding site to place the carboxylate headgroup closer to the Arg92-Arg258 pair (Fig. S22a and c-f). These movements decreased the PUFA carboxylate:Arg92 guanido group distance from the starting position to a constant distance of∼2.8Å (Fig. S22c-g) and illustrate that differences in lipid length and position of the final unsaturated bond can be accommodated by the PUFA site consistent with the lateral exit gap around the lipid tail. We also noticed that PUFAs that moved the most (ex. AA), could make new interactions with Arg258 (Fig. 22f), suggesting that plasticity in the arginine sandwich (Fig. S22h-i) may allow the channel to respond to varied PUFAs due to the enhanced electrostatic potential that surrounds this cation-cation pair^63^. As changes in the P1/M4 interface and its surrounding loops that comprise the PUFA site have diverse effects on the K_2P_ SF C-type gate^13,18,26,27,48,64,66^, such plasticity likely underlies the contrasting abilities of various PUFAs to inhibit or activate K_2P_13.1 (THIK-1). Together, these combined studies of the PUFA site lend further support to the importance of the K_2P_ P1/M4 interface in K_2P_ function and modulation^18,26,27,48,56,64^.

## Discussion

Structural characterization of K_2P_13.1 (THIK-1) reveals two functionally important elements, the flow restrictor and PUFA site, that embody the THIK subfamily and that are built upon the K_2P_ architecture shared with the TREK^18–20^, TWIK^21,22^, TALK^23^, and TASK^24^ channels. Distinct from other K_2P_s^13,18–24^, the K_2P_13.1 (THIK-1) aqueous inner cavity is divided into two chambers, the vestibule and pond, by the flow restrictor Tyr273 from the M4 helix. The hydrophilic nature of the flow restrictor, a barrier in which the component amino acids are hydrogen bonded (Figs. 1b and 2a), sets it apart from classical ion channel gates where permeation barriers comprise hydrophobic residues^61^ and provides a rationale for the observation that K_2P_13.1 (THIK-1) has one of the smallest single channel conductances among K_2P_s^5^. Movement of the hydroxyls forming this barrier enable it to affect permeant ion passage from the cytoplasmic vestibule into the pond and control ion flux (Figs. 2a and 3, Movies S1-S2). The unusual physicochemical nature of the flow restrictor provides an important example of the diverse ways that an ion channel family can be modified to affect function.

All K_2P_s, except for THIK subfamily members (Figs. S5 and S6d)^25^, have a proline at a site where there is a natural kink in the M2 transmembrane helix (Fig. S6h). The structure of the S136P mutant shows that this M2 break accommodates the activating proline mutation with minimal local helical backbone changes (Fig. 2e). This local structural similarity between wild-type and S136P channels (Fig. 2e) is consistent with ideas regarding a proline origin for nonproline kinks in transmembrane protein helices^67^ and suggests that the THIK subfamily originated from a predecessor having a proline at this site. Linkage of flow restrictor movement to subtle M2 structure changes (Figs. 2a-d), together with the rapid relaxation of the flow restrictor residue to S136P-like states in simulations (Fig. S20) suggests that the S136P mutation exploits M2 helix flexibility at the helical kink to influence Tyr273 flow restrictor conformation (Fig. S6h, Movies S1 and S2). The resulting widening of the flow restrictor barrier shows how proline substitution at this M2 site increases activity of both THIK subfamily members, K_2P_13.1 (THIK-1)^25^ and K_2P_12.1 (THIK-2)^25^, and emphasizes the role of the M2 helix in shaping K_2P_ function^13,56,68–70^. Together, our observations strongly support the idea that Tyr273 acts as a barrier that limits passage of ions from the cytoplasmic cavity into the pond and that the perturbation of this barrier is central to THIK channel function (Fig. 4j).

The K_2P_13.1 (THIK-1) PUFA site is occupied by a single chain lipid buried in the P1/M4 interface (Fig. 1c-d) and uses a signature element of the THIK subfamily, the arginine sandwich formed by the stacking of Arg92 and Arg58 guanidium groups, to interact with the PUFA carboxylate (Fig. S6c and f). This stacked cation-cation pair creates an enhanced electrostatic potential that influences interactions with oppositely charged partners^63^ and appears to enable the PUFA site to accommodate PUFAs from both ω3 and ω6 classes (Figs. 4, S21-S22). Remarkably, the THIK PUFA site structure encapsulates the essence of the TREK subfamily K_2P_ modulator pocket where synthetic small molecule activators bind^13,18,26,27^ (Figs. 1g-i). This structural coincidence of ligand binding sites shows that, as anticipated^13,18^, this interdomain modulator binding site exists in diverse K_2P_ subfamilies and can interact with natural ligands.

The K_2P_13.1 (THIK-1) PUFA appears to act as a structural co-factor, as evidenced by its co-purification with the channel (Figs. S2b and S3b-c) and the modest responses of the wild-type channel to application of a variety of PUFAs (Figs. 4d-h and S21). Notably, weakening the interactions with the PUFA carboxylate headgroup via the R92A mutation sensitizes the channel to these same lipids. K_2P_13.1 (THIK-1) R92A can be activated by linoleic acid, a PUFA that fits the observed experimental density (Figs. S2b and S3c), with a stoichiometry that matches the number of observed PUFA sites (Figs. 4i) and the response of rat K_2P_13.1 (THIK-1) to arachidonic acid^1^. Although THIK channel pharmacology remains limited^32^, it is noteworthy that the R92A change also attenuates K_2P_13.1 (THIK-1) inhibition by IBMX^53^, suggesting that the PUFA site participates in the response of THIK channels to exogenous agents. Together, these observations expand the role for the P1/M4 interface in the function of diverse K_2P_ subfamilies^18,26,27,48,56,64^ and raise the possibility that PUFAs or other endogenous factors modulate other K_2P_s^10–13^ at the equivalent interface.

SF C-type gating is the main K_2P_ gating mechanism^12,13,18,27,48–50^ and is operative in other potassium channel classes^13,71^. Notably, functional and simulation studies indicate that PUFAs such as linoleic acid activate the voltage-gated potassium channel Kv7.1 (KCNQ1) by stabilizing its SF gate through binding to a site, termed ‘Site II’, framed by the pore domain P and S6 helices^28–31^. Comparison of the Kv7.1 (KCNQ1) and K_2P_13.1 (THIK-1) pore domains shows not only that Kv and K_2P_ channel pores share a common architecture (RMSD_Cα_ = 0.257 for SF-P helix) (Fig. S23) but also that the Kv7.1 (KCNQ1) Site II coincides with the K_2P_13.1 (THIK-1) P1/M4 interface PUFA site (Figs. 4k and S23). This convergence strongly suggests that the ligand binding properties of the K_2P_ PUFA site are shared with other VGIC superfamily members and that PUFA occupation of such interdomain sites influences SF gate function. Identification of this PD interdomain site that is used by small molecule activators and PUFAs in K_2P_s^18,26,27^ and PUFAs in Kvs^28–31^ provides a framework for addressing the interplay between modulator binding and channel elements that control ion flux in the VGIC superfamily. Moreover, this common regulatory framework highlights the potential of targeting this class of interdomain interfaces for modulator development.

Some K_2P_s, including the K_2P_13.1 (THIK-1)/K_2P_12.1 (THIK-2) pair^72^, can function as heterodimers^72–76^. Although the structural consequences of K_2P_ heterodimerization remain unknown^13^, the K_2P_13.1 (THIK-1) signature features, the flow restrictor and PUFA site, are present in K_2P_12.1 (THIK-2) (Fig. S6), and are thus, likely to be important in both K_2P_12.1 (THIK-2) homodimers and K_2P_13.1 (THIK-1)/K_2P_12.1 (THIK-2) heterodimers. These elements may also contribute to THIK modulation by anesthetics^1,5^, cold temperatures^5^, and small molecule modulators^32,53^. The THIK subfamily is expressed in a wide range of tissues^1^ and is thought to affect anesthesia^51^, apoptosis^52^, and ischemic responses^17^. There is growing evidence that K_2P_13.1 (THIK-1) is important for microglial function^6–9^ in neuroinflammation and synapse development^6,8,9^. The THIK subfamily structural features defined here should aid development of agents to affect Alzheimer’s^32^, ALS^32^, and autism^6^ where such processes are thought to affect disease progression.

## Supporting information

Supplementary Figs. S1-S23, Tables S1-S2, Movie S1-S4 legends, References

Movie S1 K2P13.1 (THIK-1) structural changes.

Movie S2 Cytoplasmic view of K2P13.1 (THIK-1) structural changes.

Movie S3 Dynamics of K2P13.1 (THIK-1) Tyr273 during umbrella sampling with potassium at the flow restrictor.

Movie S4 Dynamics of K2P13.1 (THIK-1) Y273F during umbrella sampling with potassium at the flow restrictor.

## Acknowledgements

We thank J. Geo for molecular biology support, H. Khant, Y. Liu and A. Cassago at the SLAC Cryo-EM Center for help with microscope handling and data acquisition, A. Mondal for critical advice on cryo-EM data analysis and model building, J.M. Rosenberg for advice on the free energy simulations, and K. Brejc for comments on the manuscript. This work was supported by NIH grant R01-MH093603 to D.L.M.

## Author Contributions

S.R.-C., S.J., F.A.-A., F.N., M.G., and D.L.M. conceived the study and designed the experiments. S.R.-C. expressed and purified the proteins and prepared the cryo-EM samples. S.R.-C. and F.A.-A. collected and analyzed the cryo-EM data. S.R.-C. and F.A.-A. built and refined the models. S.J. collected and analyzed the two-electrode voltage clamp data. F.N. and M.G. designed and analyzed the computational studies. S.R.-C., S.J., F.N. M.G., and D.L.M. analyzed the data. M.G. and D.L.M. provided guidance and support. S.R.-C., S.J., F.A.-A., F.N., M.G., and D.L.M. wrote the paper.

## Materials and correspondence

Correspondence should be directed D.L.M. Requests for materials should be directed to D.L.M.

## Data and materials availability

Coordinates and maps are deposited in the RCSB and EMDB under accession numbers: K_2P_13.1 (THIK-1):nanodiscs 9BSN and EMD-44870, K_2P_13.1 (THIK-1):detergent 8BYI and EMD-45034, K_2P_13.1 (THIK-1) S136P:nanodiscs 9C09 and EMD-45077, K_2P_13.1 (THIK-1) S136P:detergent 9C07 EMD-45075, and K_2P_13.1 (THIK-1)S136P/Y273A:nanodiscs 9BWS and EMD-44978 and will be released immediately upon publication.

## Materials and Methods

### Expression and purification of K_2P_13.1_EM_ and mutants

The gene for a construct of the human K_2P_13.1 (THIK-1) (Uniprot Q9HB14), K_2P_13.1_EM_, spanning residues 1-350 and bearing N59Q and N65Q mutations to remove glycosylation sites in the Cap domain followed by a 3C protease cleavage site, monomeric enhanced green fluorescent protein (mEGFP), and a His_8_ tag was cloned into a modified pFastBac expression vector in which the polyhedrin promotor was replaced by a mammalian cell active CMV promotor^77^. Expression vectors for S136P and S136P/Y273A mutants were made on this background. These vectors were used for recombinant bacmid DNA generation using chemically competent DH10EmBacY (Geneva Biotech) cells. These bacmids were then used to transfect *Spodopetera frugipdera* (SF9) cells to make baculoviruses for the constructs of interest^78^.

Flasks of Expi293F cells (Gibco A14528) were grown to 2.5–3 x 10^6^ cells mL^-1^ at 37 °C supplemented with 8 % CO_2_ shaking at 120 RPM before transduction with 10 % (v/v) baculovirus stock of K_2P_13.1_EM_ or mutants. 16 – 24 h after transduction, 10 mM sodium butyrate was added to the culture to enhance protein expression^79^. Flasks were then moved to 30 °C for 24 h before harvesting the cells using centrifugation at 5,000 x g for 20 min. The pellet was gently washed with Dulbecco’s phosphate buffered saline (Gibco 14190144) to remove leftover culture media, centrifuged (3500 x g for 20 min) to recover the cell pellet, and then flash frozen with liquid nitrogen for storage at – 80°C.

A pellet from ∼5 L culture was resuspended in 250 mL hypotonic buffer (20 mM KCl, 1 mM, 10 mM Tris-HCl pH 7.8, phenylmethylsulfonyl fluoride (PMSF), 0.1 mg mL^-1^ DNase1, supplemented with Pierce protease inhibitor (Thermo Fischer Scientific) and was gently stirred on a MONO DIRECT Variomag magnetic stirrer (Thermo Fischer Scientific) at 4 °C for 30 min. The membranes were isolated by ultracentrifugation at 185,500g for 30 mins. Using a 16 G Precision Glide needle and 30 mL syringe, the membrane pellet was homogenized in 300 mL solubilization buffer (buffer S) (500 mM KCl, 50 mM Tris-HCl pH 7.8, supplemented with 1 mM PMSF, 0.1 mg mL^-1^ DNase1, 1 % (w/v) n-Dodecyl-β-D-Maltopyranoside (β-DDM), 0.2 % (w/v) Cholesteryl Hemisuccinate Tris Salt (CHS), and Pierce protease inhibitor. Protein extraction was performed by gently stirring at 4°C for 2 h and ultracentrifugation at 185,500g for 1 h to remove cell debris. The supernatant was diluted 1:1 with buffer S, supplemented with 0.1% β-DDM and 0.02% CHS, to reduce the detergent concentration and incubated for 3 h with 10 mL Talon metal affinity resin (Takara Biotech) pre-equilibrated with buffer S supplemented with 0.1% β-DDM and 0.02% CHS. Resin was then collected in an Econo-Pac chromatography column (Bio-Rad) and washed with 10 CV of buffer S supplemented with 0.5 % β-DDM and 0.1 % CHS, followed by 10 CV supplemented with 0.2 % β-DDM and 0.04 % CHS, 10 CV of buffer S supplemented with 0.1 % β-DDM, 0.02 % CHS, and finally 10 CV wash buffer (500 mM KCl, 0.05 % β-DDM, 0.01 % CHS, 50 mM Tris-HCl pH 7.8). The resin was then washed using 10 CV wash buffer supplemented with 10 mM Imidazole, and then with 10 CV of wash buffer supplemented with 30 mM Imidazole. K_2P_13.1_EM_-GFP was eluted with 2 CV wash buffer supplemented with 500 mM Imidazole and the eluent was concentrated using Amicon Ultra-15 centrifugal unit with 100 kDa cutoff to a final volume of 6 mL. 2 x Econo-Pac 10DG desalting columns (Bio-Rad) were equilibrated using 20 mL desalting buffer (200 mM KCl, 0.05 % β-DDM, 0.01 % CHS, 20 mM Tris-HCl pH 7.8) and 3 mL concentrated K_2P_13.1_EM_-GFP was applied to each desalting column and eluted using 4 mL desalting buffer before concentrating to 5 mg mL^-1^ using an Amicon Ultra-15 (100 kDa cutoff) filter.

To make nanodisc (ND) complexes, brain Extract Polar lipid (Avanti polar lipids) was prepared by dissolving in chloroform, drying under vacuum, and storing at −80°C. For reconstitution, the lipid was dissolved at 80 mM stock solution by sonication in 200 mM KCl, 0.5 % β-DDM, 0.1 % CHS, 20 mM Tris pH 7.8. K_2P_13.1_EM_ from the metal affinity chromatography step, purified MSP1E1 ^80^, and prepared lipid stock were incubated together at 1:3:300 molar ratio for 1 h in a microcentrifuge tube gently rotating at 4 °C on a Boekel 260200 Orbiton platform rotator. Bio-beads SM-2 adsorbents (Bio-rad #1528920) were prepared by washing first with methanol, followed by water and then with SEC buffer (200 mM KCl, 20 mM Tris-HCl pH 7.8). For detergent removal, 120 mg of prepared bio-beads were added per 1 mL reaction mixture and incubated for 1 h gently rotating at 4 °C. The reaction mixture was moved to another tube using a 27 G Precision Glide needle and incubated with another 120 mg aliquot of prepared bio-beads per mL rotating gently at 4 °C overnight.

200 µL anti-GFP nanobody Sepharose resin^81^ pre-equilibrated with SEC buffer was combined with 1 mL of the recovered sample and incubated in 2 mL microcentrifuge tubes gently rotating at 4 °C for 2 h. The resin was collected in a column and washed using 10 CV SEC buffer to remove unbound components including empty NDs. On-column phosphatase treatment was performed by incubating the resin with 10 mL SEC buffer supplemented with 1 mM MnCl_2_ and 4,000 units (10 μL) Lambda Protein Phosphatase (LPP) (New England BioLabs P0753L) standing at 4°C overnight. The resin was then washed using 5 CV SEC buffer and the mEGFP affinity tag was cleaved using 10 mL SEC buffer supplemented to 400 mM KCl, 1 mM EDTA, and 3C protease^82^ at a ratio of 50:1 resin volume: protease volume, treated standing at 4°C overnight. The cleaved ND sample was eluted using 10 mL SEC buffer, concentrated to 0.5 mL using an Amicon Ultra-15 (100 kDa cutoff) before purification by Size Exclusion Chromatography on Superose 6 increase 10/300 column pre-equilibrated with SEC buffer. Peaks obtained were validated by SDS-PAGE before finally concentrating to 0.6-1.0 mg mL^-1^ for Cryo-EM sample preparation. Protein concentrations were determined by absorbance ^83^ using extinction coefficients of 104,740 and 171,225 M^-1^ cm^-1^ for detergent and nanodisc samples, respectively.

For the preparation of detergent samples, hypotonic wash and solubilization were performed as described above except that both buffers were supplemented with 1 mM TCEP. The supernatant acquired after ultracentrifugation post-solubilization was incubated with the anti-GFP nanobody Sepharose resin (1 ml resin per 10g Expi293F cells) gently rotating at 4 °C for 2 h. Resin wash, on column treatment with LPP and 3C protease was performed as described above. The cleaved sample was eluted in SEC buffer supplemented with 0.05% β-DDM, 0.01 % CHS and 1 mM TCEP, concentrated to 1 mL using Amicon Ultra-15 (100 kDa cutoff) before purification by Size Exclusion Chromatography using Superdex 200 increase 10/300 GL column equilibrated with 200 mM KCl, 1 mM TCEP, 0.01% GDN, 20 mM Tris-HCl pH 7.8. Peaks obtained were validated by SDS-PAGE before final concentration using Amicon Ultra-0.5 mL (100 kDa cutoff) to 2-2.5 mg mL^-1^ for cryo-EM sample preparation.

### Cryo-EM sample preparation and Data collection

3 μL of concentrated protein sample was applied to a glow-discharged (30 s at 15 mA) grid (Quantifoil Au or Cu R1.2/1.3 300 mesh) and after a wait time of 5 s, grids were blotted once for 2.5 – 6 s (4 °C and 100 % humidity) using a FEI Vitrobot Mark IV (Thermo Fisher Scientific) and plunge-frozen in liquid ethane. The grids were screened on 200 kV Talos Arctica cryo-TEM (Thermo Fisher Scientific) equipped with K3 direct detector camera (Gatan) at the University of California, San Francisco (UCSF) EM facility. Micrographs were collected using EPU data acquisition software on FEI Titan Krios (Thermo Fisher Scientific) (SLAC National Accelerator Laboratory) at 300 keV, equipped with a K3 direct-electron detector and post-BioQuantum GIF energy filter (Gatan), set to a slit width of 20 eV. Super-resolution counting mode at a nominal magnification of x105,000 with a pixel size of 0.43 Å (physical pixel size of 0.864 Å) and a nominal defocus range between −0.8 to −2.2 μm was used (exposure time 2.5 sec) and 50 frame movie stacks were collected with a total dose of ∼50 e^-^ / Å^2^.

### Structure determination (image processing, model building and refinement)

A total of 14,254, 13,491, 13,544, 6,830, and 16,053 movie stacks were collected for K_2P_13.1_EM_:ND, K_2P_13.1_EM_:detergent, K_2P_13.1_EM_ S136P:ND, K_2P_13.1_EM_ S136P:detergent, and K_2P_13.1_EM_ S136P/Y273A:ND samples, respectively. Cryo-EM data processing was performed using cryoSPARC (version 3.2.0)^84^. Patch motion correction (cropped by a factor of 2) and patch CTF estimation (default parameters) were performed before manually curating the exposures by ice-thickness, CTF-fit resolution, total full-frame motion, and motion curvature. Selected micrographs were used for reference-free circular blob picker (diameter 80-200 Å) for particle picking followed by extraction at a box size of 288 Å. Several rounds of 2D classification were performed and further cleaning was done by categorizing particles by 3D Ab initio reconstruction and heterogeneous refinement. Finally, non-uniform and local refinement were performed to improve resolution and map quality.

The optimal sharpening for each density map was performed by phenix.auto_sharpen tool in Phenix^85,86^. A preliminary model for K_2P_13.1_EM_ was built using K_2P_3.1 (TASK1) (PDB:6RV2)^24^ as this K_2P_ is closest in sequence to K_2P_13.1 (THIK-1) and docked in the final map using phenix.dock_in_map. The model then was manually adjusted to fit the density using Coot (version 0.9.8.7)^87^ and disordered residues from the N, C terminus, and M2-M3 loop were removed. Iterative structure refinement and model building were performed using phenix.real_space_refine^88^ before adding cofactors and further refinement. The final geometry validation statistics were calculated by MolProbity^89^ and refinement statistics were generated using the comprehensive validation (cryo-EM) function in Phenix. Figures were prepared and model comparisons were performed using UCSF Chimera (version 1.15) (Pettersen et.al. Comput Chem 2004) and the Pymol package (http://www.pymol.org/pymol). Contact analysis was performed using LIGPLOT^90,91^.

### Two-electrode voltage-clamp (TEVC) electrophysiology

*Xenopus laevis* oocytes were harvested according to UCSF IACUC Protocol AN193390 and digested using collagenase (Worthington Biochemical Corporation, #LS004183, 0.8-1.0 mg/mL) in Ca^2+^-free ND96 (96 mM NaCl, 2 mM KCl, 3.8 mM MgCl_2_, 5 mM HEPES pH 7.4). Oocytes were maintained at 18 °C in ND96 (96 mM NaCl, 2 mM KCl, 1.8 mM CaCl_2_, 2 mM MgCl_2_, 5 mM HEPES pH 7.4) supplemented with antibiotics (100 units ml^-1^ penicillin, 100 µg ml^-1^ streptomycin, 50 µg ml^-1^ gentamicin) and used for experiments within one week of harvest.

K_2P_ channels were subcloned into a previously reported pGEMHE/pMO vector^48^. mRNA for oocyte injections was prepared from linearized plasmid DNA (linearized with AflII) using mMessage Machine T7 Transcription Kit (Thermo Fisher Scientific). RNA was purified using RNEasy kit (Qiagen) and stored as stocks and dilutions in RNase-free water at −80 °C. Defolliculated stage V-VI oocytes were microinjected with 0.5-38 ng mRNA and currents were recorded within 18-96 h after microinjection. For amplitude comparisons, currents were compared at the same time of expression for sample injected with equivalent amounts of mRNA. For PUFA response studies, cells expressing the equivalent amplitudes of basal current were compared.

For recordings, oocytes were impaled with borosilicate recording microelectrodes (0.3–2.0 MΩ resistance) backfilled with 3 M KCl and were subjected to constant perfusion of ND96 at a rate of 3-5 ml min^−1^. Except where otherwise indicated, recording solution was 96 mM NaCl, 2 mM KCl, 1.8 mM CaCl_2_, and 1.0 mM MgCl_2_, buffered with 5 mM HEPES, pH 7.4. Arachidonic acid (AA) (Sigma-Aldrich, A3611, CAS number: 506-32-1), Arachidonic Acid Methyl Ester (AAMe) (Sigma-Aldrich, A9298, CAS number: 2566-89-4), Eicosapentaenoic acid (EPA) (Sigma-Aldrich, E2011, CAS number: 10417-94-4), dihomo-γ-linolenic acid (DGLA) (Sigma-Aldrich, E4504, CAS number: 1783-84-2), α-linolenic acid (α-LNL) (Sigma-Aldrich, L2376, CAS number: 463-40-1), Linoleic Acid (EIC) (Sigma-Aldrich, L1376, CAS number: 60-33-3) were prepared immediately prior to use from DMSO stocks (100 mM), with final DMSO concentrations of 0.1%. For each recording, control solution (ND96) was perfused over a single oocyte until current was stable before switching to solutions containing the test compounds at various concentrations. Currents were evoked from a −80 mV holding potential followed by a 500 ms ramp from −140 mV to +50 mV. Data were recorded using a Axoclamp 900A amplifier (Molecular Devices) controlled by pCLAMP10.7 software (Molecular Devices), and digitized at 1 kHz using Digidata 1550B digitizer (Molecular Devices). Representative traces and dose response plots were generated in GraphPad Prism 9 (GraphPad Software, Boston). Dose response was fit with the Hill equation (I/I_0_=minimum+ (X^Hillslope)*(max-min)/(X^n + EC_50_^n)) to determine the half maximal effective concentration (EC_50_) and Hill coefficient ‘n’.

### Molecular dynamics and computational analysis

#### Simulation Methods

##### General setup

Simulation systems were prepared using CHARMM-GUI^92,93^. Experimental structures of K_2P_13.1 (THIK-1), S136P, and S136P/Y273A (residues 14-167 and 188-295) were embedded in a pure 1-palmitoyl-2-oleoyl-sn-glycero-3-phosphocholine (POPC) bilayer. For S136, missing residues 163-167 were transferred from the S136P structure after superposing the channels. Neutral terminal capping was used, and mutation of Y273 to F or A performed as appropriate to generate Y273F, Y273A, and S136P/Y273A structures. The system was solvated with neutralizing 0.18 M KCl in a box size of 130 x 130 x 130 Å^3^. Fatty acids (arachidonic acid ACD, di-homo-γ linoleic acid LAX, alpha-linoleic acid LNL or linoleic acid EIC) were placed in the lipid binding site as appropriate. System builds had ∼100,000 atoms. A list of all simulations is found in Table S2.

All simulations were performed using Gromacs 2020.6^94^, with the CHARMM-36m forcefield^95^ and TIP3P water. Potassium ions were manually placed in the filter, as described below. We first performed energy minimization followed by a multi-step equilibration gradually reducing restraints on protein and lipid atoms over 5 ns. For arachidonic acid, an additional initial energy minimization step was performed to ensure all double-bonds adopted the cis conformation, using small (<0.5Å) manual movement of FA carbons and a modified forcefield penalizing the trans conformation. The original forcefield was used in all subsequent steps.

A simulation timestep of 2 fs was used. Coordinates were saved every 100ps. Temperature was maintained at 303.15 K using v-rescale (time constant 1 ps) and pressure at 1 atm using a Parinello-Rahman semi-isotropic barostat^96^ (time constant 5 ps, compressibility 4.5e-5 bar^-1^). Hydrogens were constrained using the LINCS^97^ algorithm. Non-bonding interactions were cut off at 1.2 nm, with force-switching from 1.0 nm. The Particle-Mesh Ewald^98^ method was used to treat long-range electrostatic interactions. The nearest neighbor list was updated every 20 steps with a cutoff at 1.2 nm and a Verlet cutoff scheme.

##### Unbiased simulations

For unbiased molecular dynamics simulations, four K^+^ were placed in the filter at locations observed in the structures (S1, S2, S3, S4). For each build system, we performed three independent repeats starting from the same energy-minimized structure, followed by equilibration and then a 200ns production run.

##### Umbrella sampling simulations

The potential of mean force (PMF) for a K^+^ entering the channel from the cytoplasm was determined along the z-axis through the central pore, parallel to the membrane normal. Specifically, the reaction coordinate was taken as the z distance from the K^+^ to the center-of-mass of the CA atoms of residues 138-144 and 272-278 (corresponding to the location of the flow restrictor). Windows were created at 1 Å intervals between +7 and −13 Å, and then at −14.5, −16, −17.5, −19 and −20.5 Å, for a total of 26 windows. The reaction coordinate value was restrained with a harmonic potential with force constant 1000 kJ mol^-1^ nm^-2^. An additional flat bottom restraint with 15 Å radius was applied in the xy-plane to prevent the ion from moving away from the protein as the windows approach the bulk. Reaction coordinate values were saved every 1000 steps.

For all umbrella windows, three K^+^ were restrained in the S0, S2 and S4 sites (based on the most common ion configuration observed in our unbiased simulations) using a flat-bottom potential in the z-direction with width 2 Å and force constant 1000 kJ mol^-1^ nm^-2^, centered on the center-of-mass of the O atoms of residues (114, 115, 241, 242); (112, 113, 239, 240); and (110, 111, 237, 238), respectively.

Equilibration was performed separately for each umbrella window with the umbrella restraints, before a production run of 100ns. For K_2P_13.1(THIK-1) and Y273F, selected windows when the potassium ion is passing the flow restrictor (z=-1 to 2 Å) were run in duplicate.

##### Analysis

Analysis was performed using MDAnalysis^99,100^. To investigate the behavior and effect of bound FA, protein snapshots were first aligned to the respective initial experimental structure using the CA atoms of the transmembrane portion (resid 14-42, 96-162, and 188-295) and the RMSD of the FA (C atoms) or protein residues comprising the FA binding site (CA of resid 97-110, 254-270) relative to the initial structure measured. We excluded the first 50 ns in all calculations to allow for equilibration.

PMFs were obtained from umbrella sampling simulations using the Grossfield wham analysis tool^101^. Convergence was assessed by running wham on blocks of 20ns, starting at 5ns intervals; times before the profile settles is discarded as equilibration. Profiles were aligned by setting ‘bulk’ to 0 kcal mol^-1^, using the average value between −15 and −20 Å.

