## Supplementary Figs. S1-S23, Tables S1-S2, Movie S1-S4 legends, References for "Structure of the human K_2P_13.1(THIK-1) channel reveals a novel hydrophilic pore restriction and lipid cofactor site"

26 June 2024

[6]Molecular Biophysics and Integrated Bio-imaging Division
Lawrence Berkeley National Laboratory, Berkeley, CA 94720 USA

### Current address: Department of Physiology, David Geffen School of Medicine at UCLA, Los Angeles, CA

Keywords: $K_{2P}$ channel, lipid nanodiscs, cryo-electronmicroscopy electrophysiology, polyunsaturated fatty acid (PUFA)

**Figure S1** **Roy-Chowdhury, *et al.***

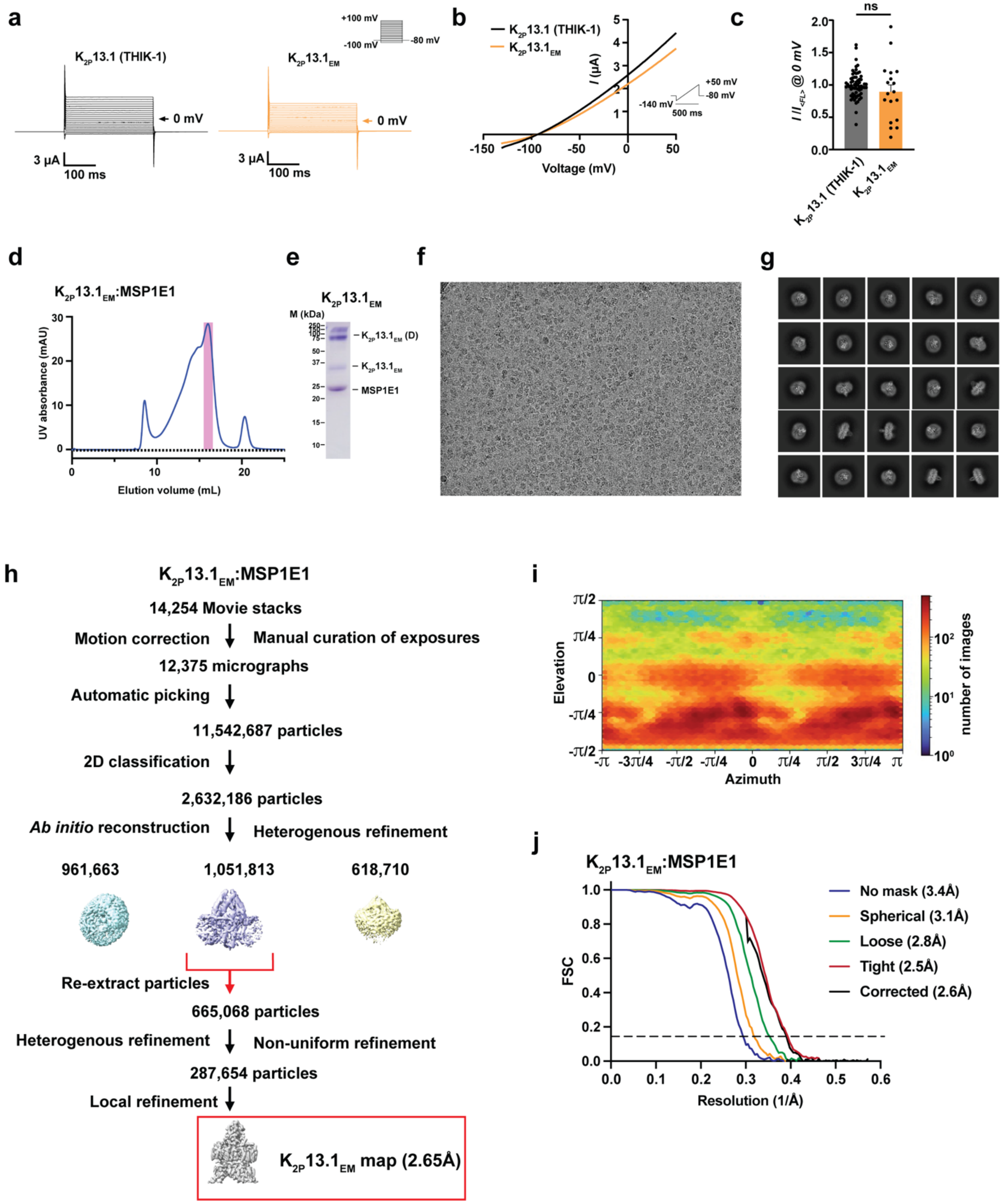

**Figure S1 Functional characterization of $K_{2P}13.1_{EM}$ and Cryo-EM analysis in MSP1E1 nanodiscs. a,** Exemplar Two Electrode Voltage Clamp (TEVC) recordings of $K_{2P}13.1$ (THIK-1) (grey) and $K_{2P}13.1_{EM}$ (orange) in *Xenopus* oocytes. 0 mV trace is indicated. Inset shows protocol. **b,** Representative TEVC current-voltage responses for $K_{2P}13.1$ (THIK-1) (black) and $K_{2P}13.1_{EM}$ (orange). Inset shows protocol. **c,** Average currents $I/I_{<FL>}$ at 0 mV where $I_{<FL>}$ = average currents for full length $K_{2P}13.1$ (THIK-1). n.s. $p > 0.05$. Statistical analysis was performed using Kruskal-Wallis test (nonparametric ANOVA) followed by Dunn's multiple comparisons test. **d,** Exemplar SEC (Superose 6 Increase 10/300 GL) for $K_{2P}13.1_{EM}$:MSP1E1 nanodiscs. **e,** peak fraction SDS-PAGE. **f,** electromicrograph (~105,000x magnification), and **g,** 2D class averages. **h,** Workflow for cryo-EM data processing for the $K_{2P}13.1_{EM}$:MSP1E1 nanodisc complex in cryoSPARC-3.2[1]. Red arrow indicates the class of particles extracted without Fourier cropping after the initial cleanup. These were further subjected to heterogeneous, non-uniform, and local refinement jobs that resulted in the final map at 2.65Å (red box). **i,** Particle distribution plot and **j,** gold-standard Fourier Shell Correlation (FSC) curves for the $K_{2P}13.1_{EM}$:MSP1E1 nanodisc complex.

**Figure S2** **Roy-Chowdhury, *et al.***

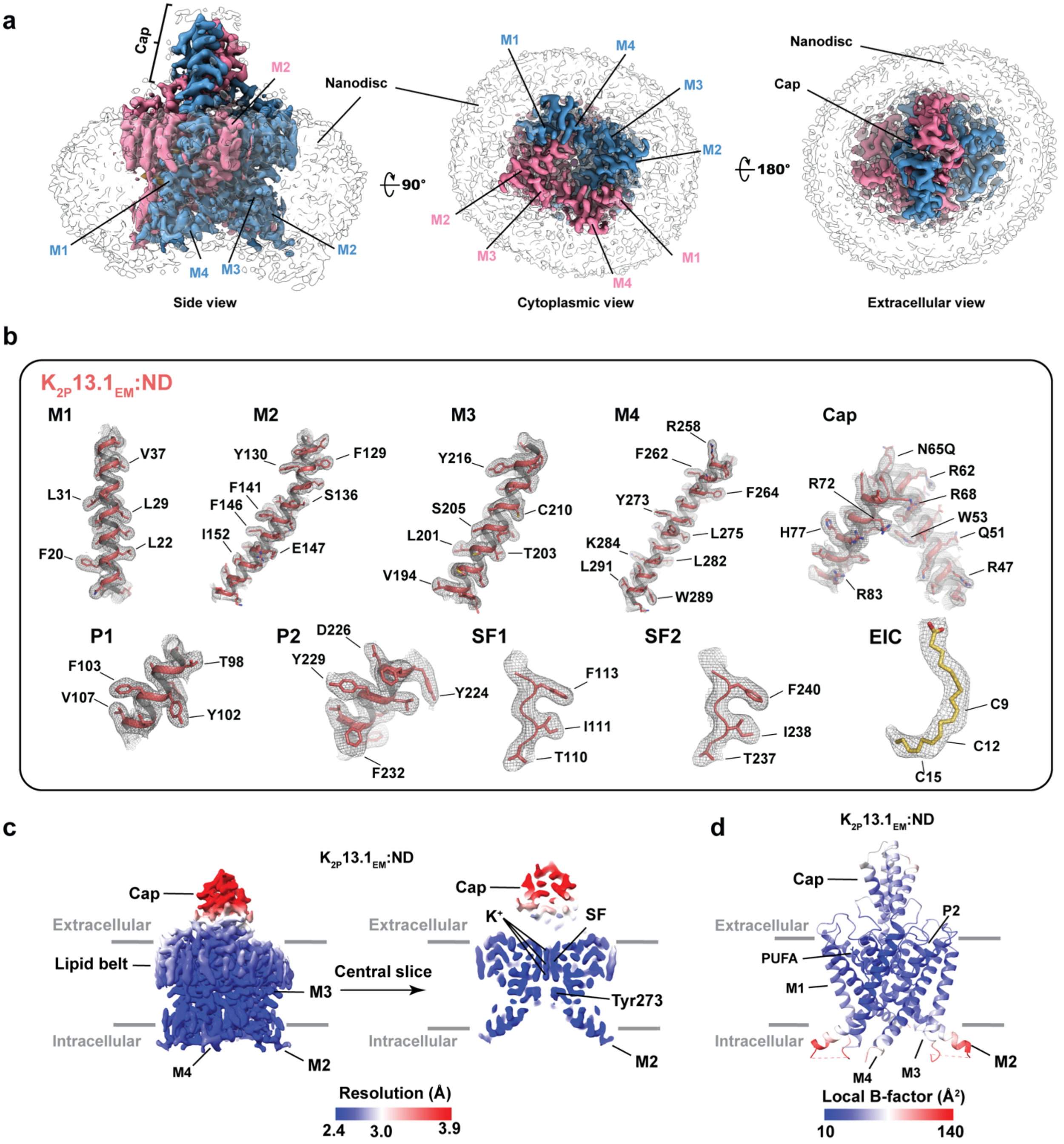


**Figure S2 $K_{2P}13.1_{EM}$:MSP1E1 nanodisc (ND) cryo-EM map and model quality. a,** Cryo-EM map of the $K_{2P}13.1_{EM}$:ND complex. $K_{2P}13.1_{EM}$ and associated lipids are colored magenta and turquoise. Locations of select channel elements are indicated. Nanodisc density is transparent. **b,** Cryo-EM maps for indicated $K_{2P}13.1_{EM}$ elements. Select residues are indicated. Channel elements are deep salmon. Linoleic acid (EIC) is yelloworange. Maps are rendered at 4-5σ.

**c,** $K_{2P}13.1_{EM}$:ND local resolution. Select channel elements and lipid belt are labeled.
**d,** $K_{2P}13.1_{EM}$:ND local B-factor.

**Figure S3** **Roy-Chowdhury, *et al.***

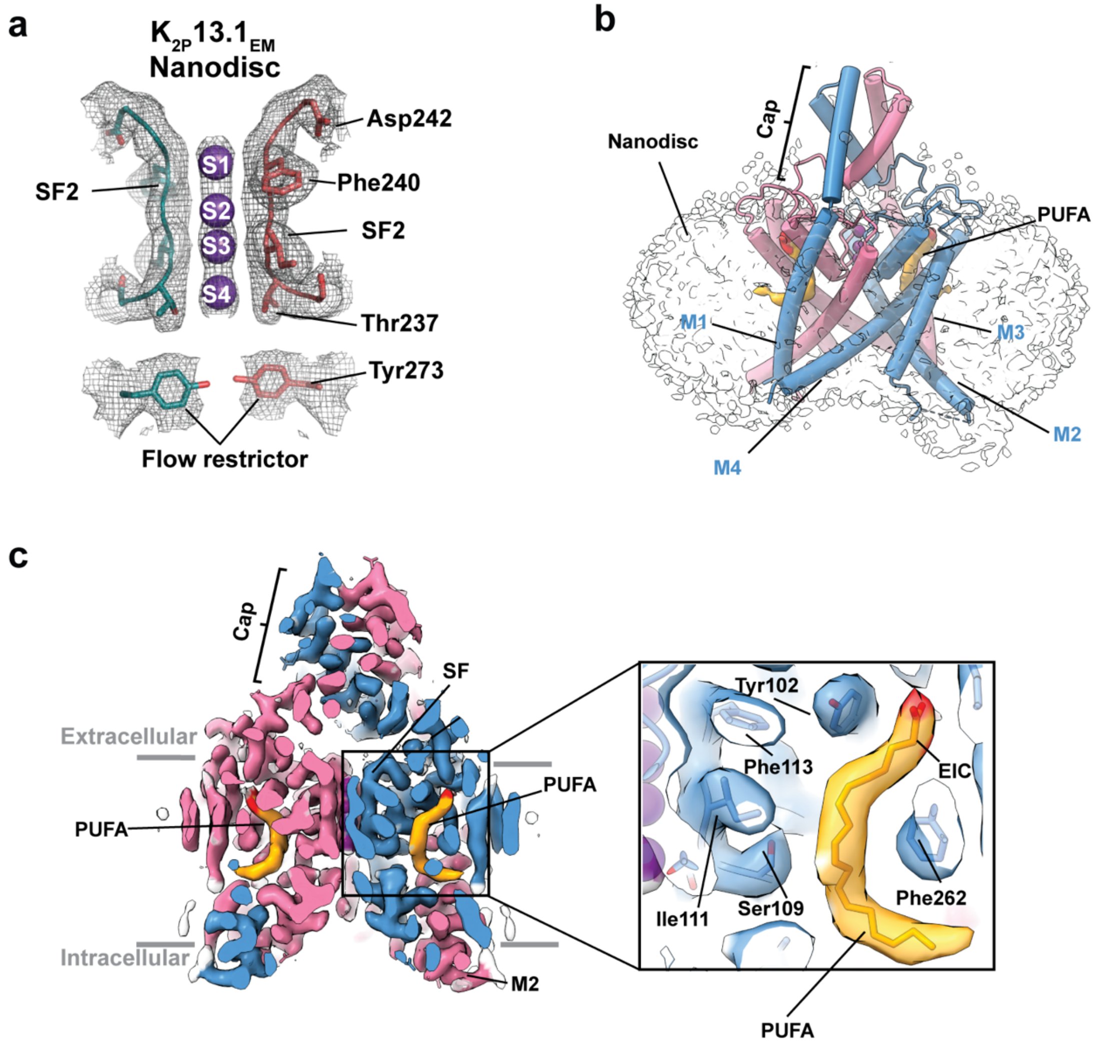


**Figure S3 $K_{2P}13.1_{EM}$:ND cryo-EM maps of key features. a,** Selectivity filter and flow restrictor densities (6σ) for $K_{2P}13.1_{EM}$:ND (deep teal and salmon). Select residues are labeled. Potassium ions are shown as purple spheres. **b,** Cyro-EM map of the $K_{2P}13.1_{EM}$:ND complex showing the channel cartoon (magenta and turquoise) and location of PUFA density (red and yelloworange). Turquoise subunit transmembrane helices are indicated. Nanodisc density is transparent. **c,** Slice through the $K_{2P}13.1_{EM}$:ND complex cyro-EM map (7σ) showing location of PUFA density (red and yelloworange). Inset shows local density. PUFA site shows linoleic acid (EIC). Grey bars denote the membrane.

**Figure S4** **Roy-Chowdhury, *et al.***

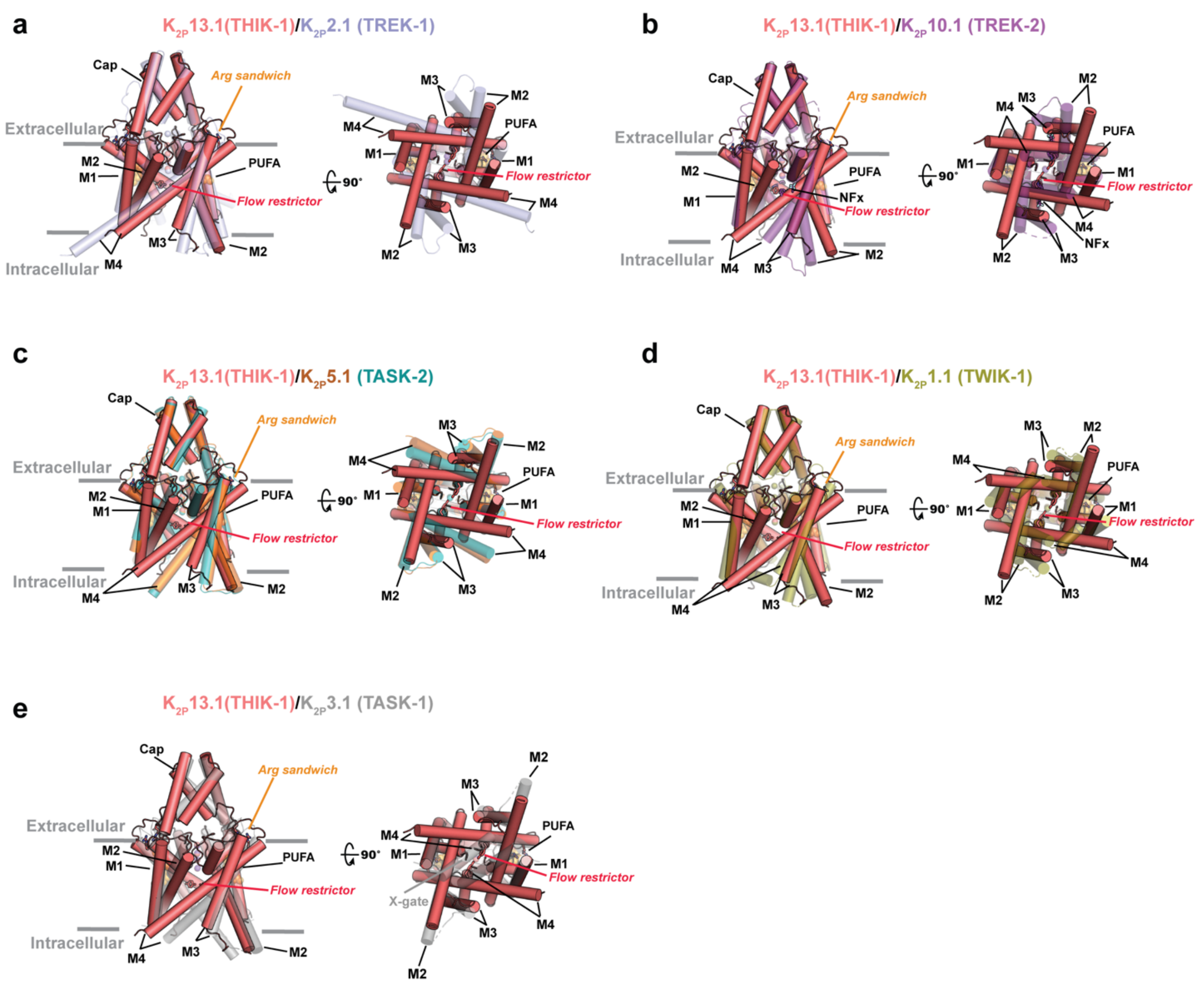


**Figure S4 $K_{2P}$13.1 (THIK-1) structure comparisons.** Superpositions of $K_{2P}$13.1 (THIK-1) (salmon) (PDB:9BSN) with: **a,** $K_{2P}$2.1 (TREK-1):ML335 (light blue) (PDB:6CQ6)[2] ($RMSD_{C\alpha}$ = 1.1613Å) **b,** $K_{2P}$10.1 (TREK-2):norfluoxetine (Nfx) (hot pink) (PDB:4XDK)[3] ($RMSD_{C\alpha}$ = 1.580Å). **c,** $K_{2P}$5.1 (TASK-2) pH 6.5 (orange) (PDB:6WLV)[4] ($RMSD_{C\alpha}$ = 1.816) and pH 8.5 (teal) (PDB:6WM0) ($RMSD_{C\alpha}$ = 1.639Å)[4]. **d,** $K_{2P}$1.1 (TWIK-1) (deep olive) (PDB:3UKM)[5] ($RMSD_{C\alpha}$ = 2.277Å). **e,** $K_{2P}$3.1 (TASK-1) (grey) (PDB:6RV2)[6] ($RMSD_{C\alpha}$ = 1.323Å). For simplicity, only one $K_{2P}$13.1 (THIK-1) Arg sandwich position and PUFA site are labeled. Site of the $K_{2P}$3.1 (TASK-1) X-gate is labeled in '**e**'. Each panel shows side (left) and cytoplasmic (right) views. Grey bars denote the membrane.

**Figure S5** **Roy-Chowdhury, *et al.***

```
                                      M1   24  27 28  31                          Cap α1
H. sapiens               1   MAGR-GFS--WGPGHLNEDNARFLLLAALIVLYLLGGAAVFSALELAHERQAKQRWEERLANFSRGHNLS   67
R. ferrumequinum         1   MAGR-GCG--CGSGHLNEDNARFLLLAALIVLYLLGGAAVFSALELAHELQAKQRWEERLANFSRRHNLS   67
C. lupus familiaris      1   MAGR-GCGCSCGPGHLNEDNARFLLLAALIVLYLLGGAAVFSALELAHERQAKQRWEERLANFSRRHNLS   69
O. orca                  1   MAGR-GCGCSCGPGHLNEDNARFLLLAALIVLYLLGGAAVFSALELAHERQAKQRWEERLANFSRRHNLS   69
R. norvegicus            1   MAGR-GCS--CSPGHLNEDNARFLLLAGLILLYLLGGAAVFSALELAQELQAKQRWEERLANFSRGHNLS   67
M. musculus              1   MAGR-GCG--CSPGHLNEDNARFLLLAGLILLYLLGGAAVFSALELAQELQAKQRWEERLANFSRGHNLS   67
B. taurus                1   MAGR-GCGGDCGPGRANEDNARFLLLGALLALYLLGGAAVFSALERPHERQARQRWEERLAGFGRRHNLS   69
G. gallus                1   MACG-GCCC-SGAGRLNADNARFLLLAVLIALYMLCGAAVFSAIELPGERRAKQRWEERLQNFTRRHNLS   68
X. tropicalis            1   MACR-GCCC-CGFGHINEDNGRFLLLAILIILYLLCGAAVFSAIEQPKEKEAKERWQQKRDTFSQKYNLN   68
D. rerio                 1   MACRSGCCC-NSIGSFNEDNARFLMLALLIIIYLLCGAAVFSALEQPKEKLAKERWAQRIEQFTNKYNLS   69

                                Cap α2            92       101 102 P1 105 108 109 111 SF1          M2     136
H. sapiens              68   RDELRGFLRHYEEATRAGIRVDNVRPRWDFTGAFYFVGTVVSTIGFGMTTPATVGGKIFLIFYGLVGCSS   137
R. ferrumequinum        68   RDELRGFLRHYEEAAKAGIRVDNTRQRWDFTGAFYFVGTVVSTIGFGMTTPATVGGKTFLILYGLIGCAS   137
C. lupus familiaris     70   RDELRGFLRHYEEATEAGIRVDSARPRWDFTGAFYFVGTVVSTIGFGMTTPATVGGKIFLIFYGLIGCAS   139
O. orca                 70   RDELRGFLRHYEEATEAGIRVDNVRLRWDFTGAFYFVGTVVSTIGFGMTTPATVEGKLFLIFYGLLGCSS   139
R. norvegicus           68   REELRGFLRHYEEATKAGIRMDSVRPRWDFTGAFYFVGTVVSTIGFGMTTPATTGGKIFLIFYGLIGCAS   137
M. musculus             68   REELRGFLRHYEEATRAGIRMDSVRPRWDFTGAFYFVGTVVSTIGFGMTTPATTGGKIFLIFYGLIGCAS   137
B. taurus               70   RDALRGFLREYEEASVAGIRVDDIRPRWDFPGAFYFVGTVVSTIGFGMATPATVGGKVFLIGYGLLGCSS   139
G. gallus               69   RAELRDFLREYEEASVAGIRVDDIRPRWDFTGAFYFVGTVVSTIGFGMTTPATVGGKIFLIFYGLIGCAG   138
X. tropicalis           69   VSDLRNFLRDYEEANVAGIRVDSIRPRWDFTGAFYFVGTVVSTIGFGMTTPVTIAGKIFLIFYGLIGCAA   138
D. rerio                70   QDDLKNFLRNYEEANVAGIRVDATRARWDFAGAFYFVGTVVSTIGFGMTTPVTIAGKIFLIFYGLIGCAA   139

                             138 141 143  M2                                                M3
H. sapiens             138   TILFFNLFLERLITIIAYIMKSCHQRQLRRRGALPQESLKDAG----QCEVDSLAGWKPSVYYVMLILCT   203
R. ferrumequinum       138   TILFFNLFLERLITVIAYIMKSCHQQQLRSQRALPQEGLKNLG----KREADGALGWKPSVYYVMLILCV   203
C. lupus familiaris    140   TILFFNLFLERLITVIAYIMKSCHQRQLQRRGALPPESLKNPG----RCEVDGLAGWKPSVYYVMLILCV   205
O. orca                140   TILFFNLFLERLITVIACIMKSCHQRQLQRRGALPQQSLKNPG----KGEADSSAGWKPSVYYVMLILCM   205
R. norvegicus          138   TILFFNLFLERLITVIAYVMRTCHQQQLRRRGTVARDNGKAPR----KGEADSLAGWKPSVYYVMLILCL   203
M. musculus            138   TILFFNLFLERLITVIACVMRSCHQQQLRRRGAVTQDNMKAPE----KGEADSLTGWKPSVYYVMLILCL   203
B. taurus              140   TLLFFNLFLERLITGIAHVTRWCGARRPGRAGR---RARSPG--------RAAAAGKPSVRVVLLALGA   198
G. gallus              139   TILFFNLFLERLITVIAYVMKSCHERQLRRKGVLPHDGRRGSG----TSEADSLAGWKPSVYYVMLILCI   204
X. tropicalis          139   TILFFNLFLERLITVAIYIMKSFHERQLKRKGGLPPETRRGSG----HSEVDSLAGWKPSVYYVMLILCV   204
D. rerio               140   TILFFNLFLERVITVIAFVLKFCHERRESRKAGPTQ-NCRRPSTDNRDRRTDSLAGWKPSVYCVMLILGV   208

                                M3                 P2             SF2   246       M4 258 261 262 265 266 273
H. sapiens             204   ASILISCCASAMYTPIEGWSYFDSLYFCFVAFSTIGFGDLVSSQNAHYESQGLYRFANFVFILMGVCCIY   273
R. ferrumequinum       204   ASVLISCCASAMYTSIEGWSYFDSLYFCFVAFSTIGFGDLVSSQNSQYKSQGLYRFANFAFILMGVCCIY   273
C. lupus familiaris    206   ASLLISCCASAMYTSIEGWSYFDSLYFCFVAFSTIGFGDLVSSQHAQYDRQGLYRFANFVFILMGVCCIY   275
O. orca                206   ASLLISCCASAMYTSVEGWSYFDALYFCFVAFSTIGFGDLVSSQNARYASQGLYRFANFVFILMGVCCIY   275
R. norvegicus          204   ASVAISCGASALYTTMEGWSYFDSVYFCFVAFSTIGFGDLVSSQNAQYENEGLYRFVNFFLILMGVCCIY   273
M. musculus            204   ASVAISCGASALYTTMEGWSYFDSVYFCFVAFSTIGFGDLVSSQNAQYESQGLYRFFNFFLILMGVCCIY   273
B. taurus              199   ASLALCCGAAALYAAAEGWGYLDALYFCFVAFSTIGFGDLVSGQQARYDGQGLYRVANFVLLLSGVCCLY   268
G. gallus              205   ASLIISCCASAMTYPIEGWSYFDSLYFCFVAFSTIGFGDLVSSQNTKYESQGLYRFGNFVFILMGVCCIY   274
X. tropicalis          205   ASIIISCCASAMFSPIEGWGYFDSLYFCFVAFSTIGFGDMVSSQKAKYENQGLYRFGNFIFILMGVCCIY   274
D. rerio               209   AAILVSCCASMAYSAAEGWDYLDALYFCFVAFSTIGFGDMVSNQREIYEAQVAYRVGNFLFILTGVCCIY   278

                             277  M4
H. sapiens             274   SLFNVISILIKQSLNWILRKMDSGCCPQCQRGLLRSRRNVVMPGS-VRNRCNISIETDGVAESDTDGRRL   342
R. ferrumequinum       274   SLFNVISILIKQSMNWILRKMDGGCCQQCQSGLSRSRRNVIMPGN-VQNRCNISIETDGVMDSDTDGRRP   342
C. lupus familiaris    276   SLFNVISILIKQSVNWILRKMEGGCCQQCRRRLGRSRRNVVMPGNSVQNRCNISIDTDGVMDSDTDGRRL   345
O. orca                276   SLFNVISILIKQSVNWILRKMDGGCCQQCQRRLLQSRRNVVMPGT-VQNRCDISIETDGVIESDTDGRRL   344
R. norvegicus          274   SMFNVISILIKQTVNWILRKLDSGCFPQCQRGLLRSRRNVVMPGN-IRNRCNISIERDGVMESDTDGRRL   342
M. musculus            274   SLFNVISILIKQTVNWILRKLDSGCFPPCQRGLLRSRRNVVMPGN-IRNRCNISIERDGVMESDTDGRRL   342
B. taurus              267   SLFNVVSILVRQAVNWALGRAAGAGCRRGRGRGGPPRRNAVGPGR-VRSGARGAAEADGS----SGARRL   333
G. gallus              275   SLFNVISIVIKQSINWILKKLDCRCCHRCQRKLFRPKRNAVMPGN-VRGRRNISIETDIVNESDTDGRRL   343
X. tropicalis          275   SLFNVISIVIKQCVNWILRKLECRCCQRCQRKLFRPKRNAVMPGN-VRTRRNISIETDGVYESETDGRRM   343
D. rerio               279   SLFNVISIVIKQVLNWLLRRLETPCHCP-GRGNRHPRRNAVVPGH-MRSRRENSIETDAVNDSEADGRRM   346

H. sapiens             343   SGEMISMKDLLAANKASLAILQKQLSEMANGCPHQTSTLARDNEFSGGVGAFAIMNNRLAETSGDR   408
R. ferrumequinum       343   SGEMISMKDFLAANKVALAILQKQLSETANGCPHQTSTLSRDNEFSGGVGAFAIMNNRLAETSGDR   408
C. lupus familiaris    346   SGEMISMKDFLAANKVSLAILQKQLSETANGGPPQASVLSRDSEFSGGVGAFAIMNNRLAETSGDR   411
O. orca                345   SGEMISMKDFLAANKVSLAILQKQLSETANGCPHQTSTLSLDDEFSGGIGAFAVMNNRLAETSGDR   410
R. norvegicus          343   SGEMISMKD---TNKVSLAILQKQLSEMANGGPHQTSTSSRDDEFSGGVGAFAVMNNRLAETSGDR   405
M. musculus            343   SGEMISMKD---TNKVSLAILQKQLSEMANGGPHQNSASSRDDEFSGGVGAFAVMNNRLAETSGDR   405
B. taurus              334   SGEMVSMKDLLAANKASLAALQKQLSEAASGRPRRASTPPADDEFSGGVGAFAVMNGRLAETSAGS   399
G. gallus              344   SGEMISMKDFLASNKVSLAIMQKQLSETANGYPRQMS--SKQNGFSGGVGALAIMNNRLAETSVDR   407
X. tropicalis          344   SGEMISMKDFLAANKVSLAIMQKQLSETANGCPRQIGTSSRHNGFSGGIGALAIMNNRLAERSVDR   409
D. rerio               347   SGEMISMKDFLAANKVNLAIMQKQLSEMANGHPRQSGSSSRHNEFSGGVGALGIMNNRLAETSVDR   412
```

**Figure S5 $K_{2P}$13.1 (THIK-1) sequence conservation.** Sequence alignment of *H. sapiens* $K_{2P}$13.1 (THIK-1) (NP_071337.2) with homologs from the greater horseshoe bat (*R. ferrumequinum*) (XP_032962921.1), domestic dog (*C. lupus familiaris*) (XP_038449728.1), killer whale (*O. orca*) (XP_033274470.2), rat (*R. norvegicus*) (NP_071629.2), mouse (*M. musculus*) (NP_001157898.1), cattle (*B. tarus*) (XP_024853931.1), chicken (*G. gallus*) (XP_001235376.1), Western clawed frog (*X. tropicalis*) (XP_002933246.1), and zebrafish (*D. rerio*) (NP_001020654.1). PUFA site (orange), arginine sandwich (light blue), asparagine ring (green), and flow restrictor (red) sites indicated. Purple helix indicates M2 extension seen in detergent structures. Red dashed lines indicate residues lacking cryo-EM density. Grey dashed line indicates residues absent from the $K_{2P}13.1_{EM}$ construct.

**Figure S6** **Roy-Chowdhury, *et al.***

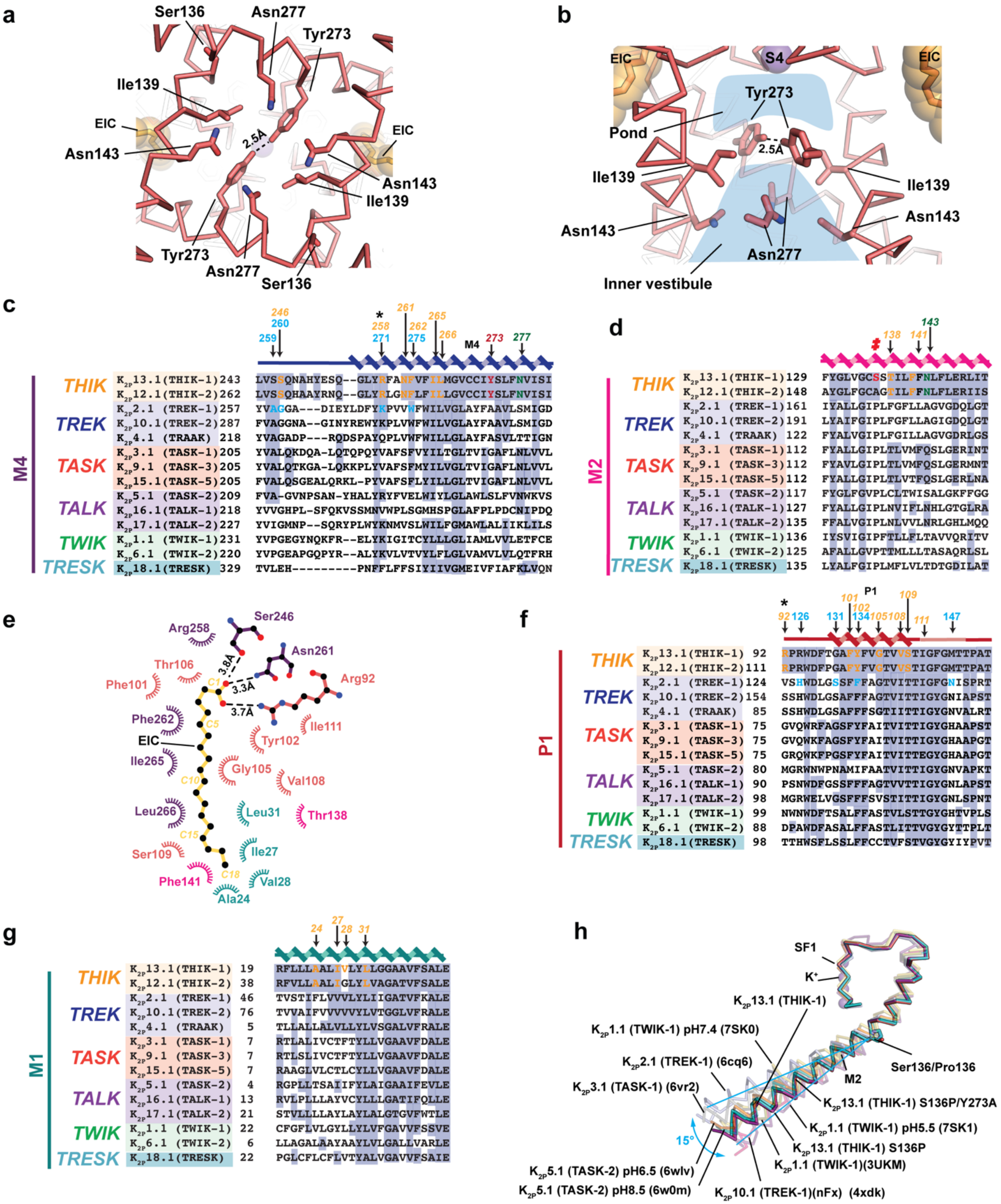


**Figure S6 $K_{2P}$13.1 (THIK-1) structural and sequence features. a,** Cytoplasmic view of the $K_{2P}$13.1 (THIK-1) Asn ring (Asn134 and Asn227) and flow restrictor barrier (Ile139 and Tyr273).

Ser136 is also shown. Linoleic acid (EIC) is shown in space filling. **b,** Lateral view of the $K_{2P}$13.1 (THIK-1) central cavity. Asn ring (Asn134 and Asn227) and flow restrictor barrier (Ile139 and Tyr273) are labeled. EIC and S4 potassium ion (violet) are shown in space filling. Pond and inner vestibule regions are shown in blue. **c,** M4 sequence comparison. Labels indicate: $K_{2P}$13.1 (THIK-1) EIC contact residues (orange), flow restrictor (red), Asn ring (green), and $K_{2P}$2.1 (TREK-1) $K_{2P}$ modulator pocket residues that contact the ML335 activator (light blue)[2]. Black asterisk denotes Arg sandwich residue. **d,** M2 sequence comparison. Labels indicate $K_{2P}$13.1 (THIK-1) EIC contact residues (orange) and Asn ring (green). Red ‡ indicates S136P mutation site. **e,** PUFA site showing hydrogen bond and salt bridge interactions (dashed lines) and van der Waals contacts ≤ 5Å. Colors indicate residues from the M1 (teal), M2 (magenta), P1 (salmon), and M4 (violet) helices. EIC is yelloworange and black. **f,** P1 sequence comparison. Labels indicate $K_{2P}$13.1 (THIK-1) EIC contact residues (orange) and $K_{2P}$2.1 (TREK-1) $K_{2P}$ modulator pocket residues that contact the ML335 activator (light blue)[2]. Black asterisk denotes Arg sandwich residue. **g,** M1 sequence comparison. Labels indicate $K_{2P}$13.1 (THIK-1) EIC contact residues (orange). **h,** Superposition of the SF1-M2 regions of $K_{2P}$13.1 (THIK-1) (salmon), $K_{2P}$13.1 (THIK-1) S136P (cyan), and $K_{2P}$13.1 (THIK-1) S136P/Y273A (magenta) with $K_{2P}$2.1 (TREK-1):ML335 (light blue) (PDB:6CQ6)[2], $K_{2P}$10.1 (TREK-2):norfluoxetine (Nfx) (hot pink) (PDB:4XDK)[3], $K_{2P}$5.1 (TASK-2) pH 6.5 (orange) (PDB:6WLV)[4] and pH 8.5 (teal) (PDB:6WM0)[4], $K_{2P}$1.1 (TWIK-1) (deep olive) (PDB:3UKM)[5], pH 7.4 (7SK0)[7], and pH 5.5 (7SK1)[7], and $K_{2P}$3.1 (TASK-1) (grey) (PDB:6RV2)[6]. Cyan lines show the range of M2 movements indicated by the 15° double headed arc. Potassium ions are shown as purple spheres. GenBank Sequences in **c, d, f,** and **g** are for human: $K_{2P}$13.1 (THIK-1) 16306555; $K_{2P}$13.2 (THIK-2) 11545761; $K_{2P}$2.1 (TREK-1) 14589851; $K_{2P}$10.1 (TREK-2) 20143944; $K_{2P}$4.1 (TRAAK) 15718767; $K_{2P}$3.1 (TASK-1) 4504849; $K_{2P}$9.1 (TASK-3) 542133161; $K_{2P}$15.1 (TASK-5) 333440483; $K_{2P}$5.1 (TASK-2) 333440483; $K_{2P}$16.1 (TALK-1) 14149764; $K_{2P}$17.1 (TALK-2) 17025230; $K_{2P}$1.1 (TWIK-1) 4504847; $K_{2P}$6.1 (TWIK-2) 4758624; and $K_{2P}$18.1 (TRESK) 32469495.

**Figure S7** **Roy-Chowdhury, *et al.***

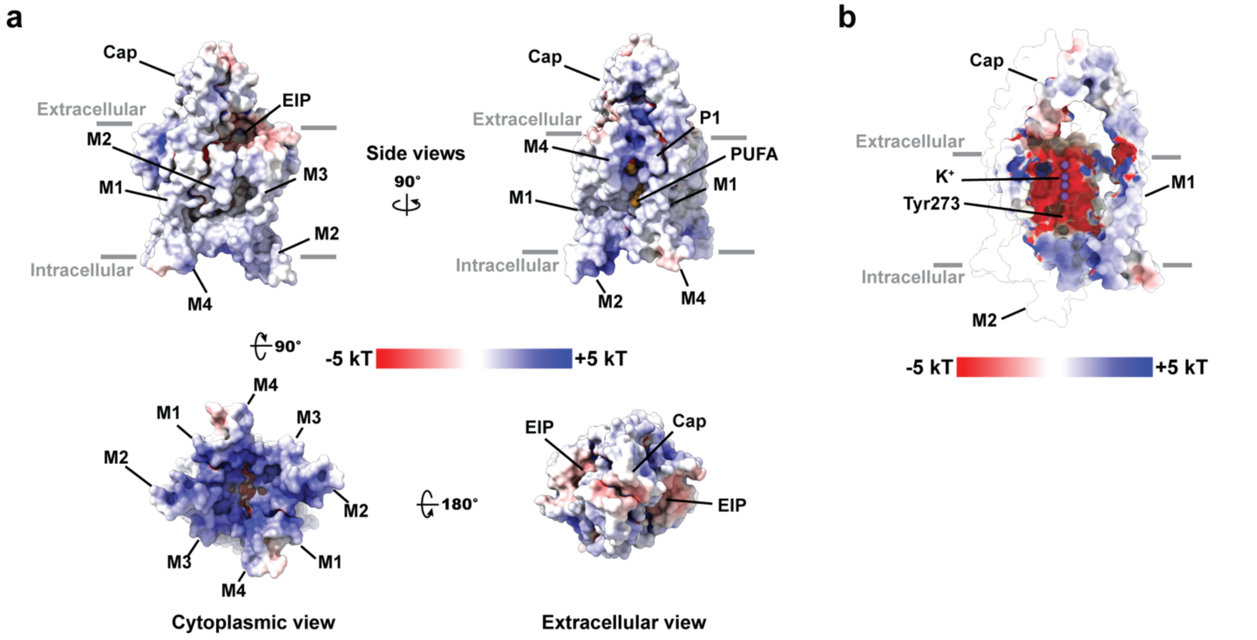


**Figure S7 $K_{2P}$13.$1_{EM}$ electrostatic surface potentials. a,** Electrostatic surface potentials calculated using APBS[8]. Side, cytoplasmic, and extracellular views are shown. **b,** Slice through the center of $K_{2P}$13.$1_{EM}$ showing the inner cavity. Selectivity filter ions are shown as purple spheres. Location of Tyr273 is indicated. One subunit is shown as transparent. Select channel elements are labeled. Grey bars denote the membrane.

**Figure S8** **Roy-Chowdhury, *et al.***

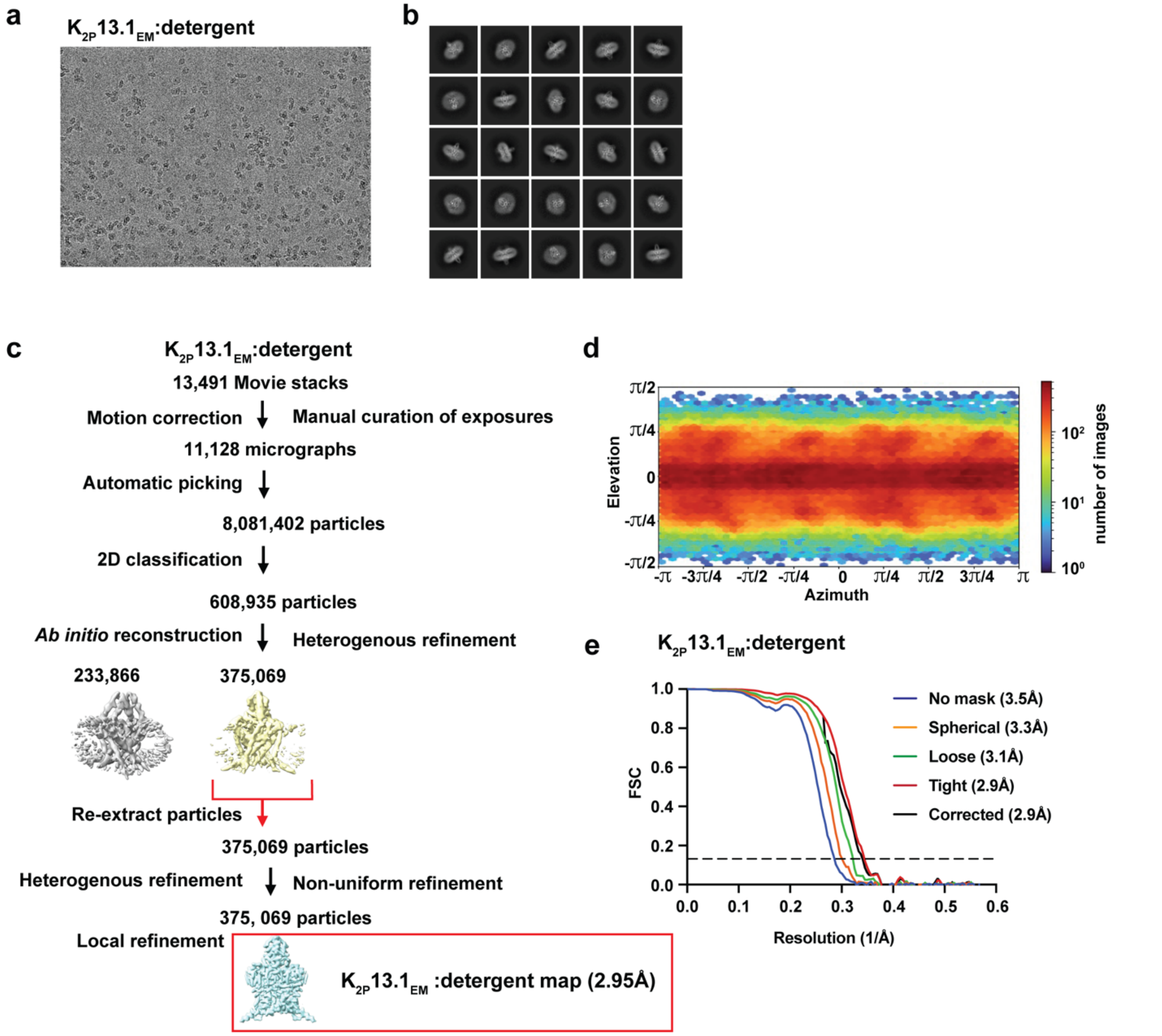


**Figure S8 Cryo-EM analysis of K$_{2P}$13.1$_{EM}$ DDM:CHS:GDN detergent micelles.** Exemplar **a,** electromicrograph (~105,000x magnification), and **b,** 2D class averages. **c,** Workflow for cyro-EM data processing for the K$_{2P}$13.1$_{EM}$:DDM:CHS:GND detergent micelle complex in cryoSPARC-3.2[1]. Red arrow indicates the class of particles extracted without Fourier cropping after the initial cleanup. These were further subjected to heterogeneous, non-uniform, and local refinement jobs that resulted in the final map at 2.95Å. (red box). **d,** Particle distribution plot and **e,** gold-standard Fourier Shell Correlation (FSC) curves for the K$_{2P}$13.1$_{EM}$:DDM:CHS:GND detergent micelle complex.

**Figure S9** **Roy-Chowdhury, *et al.***

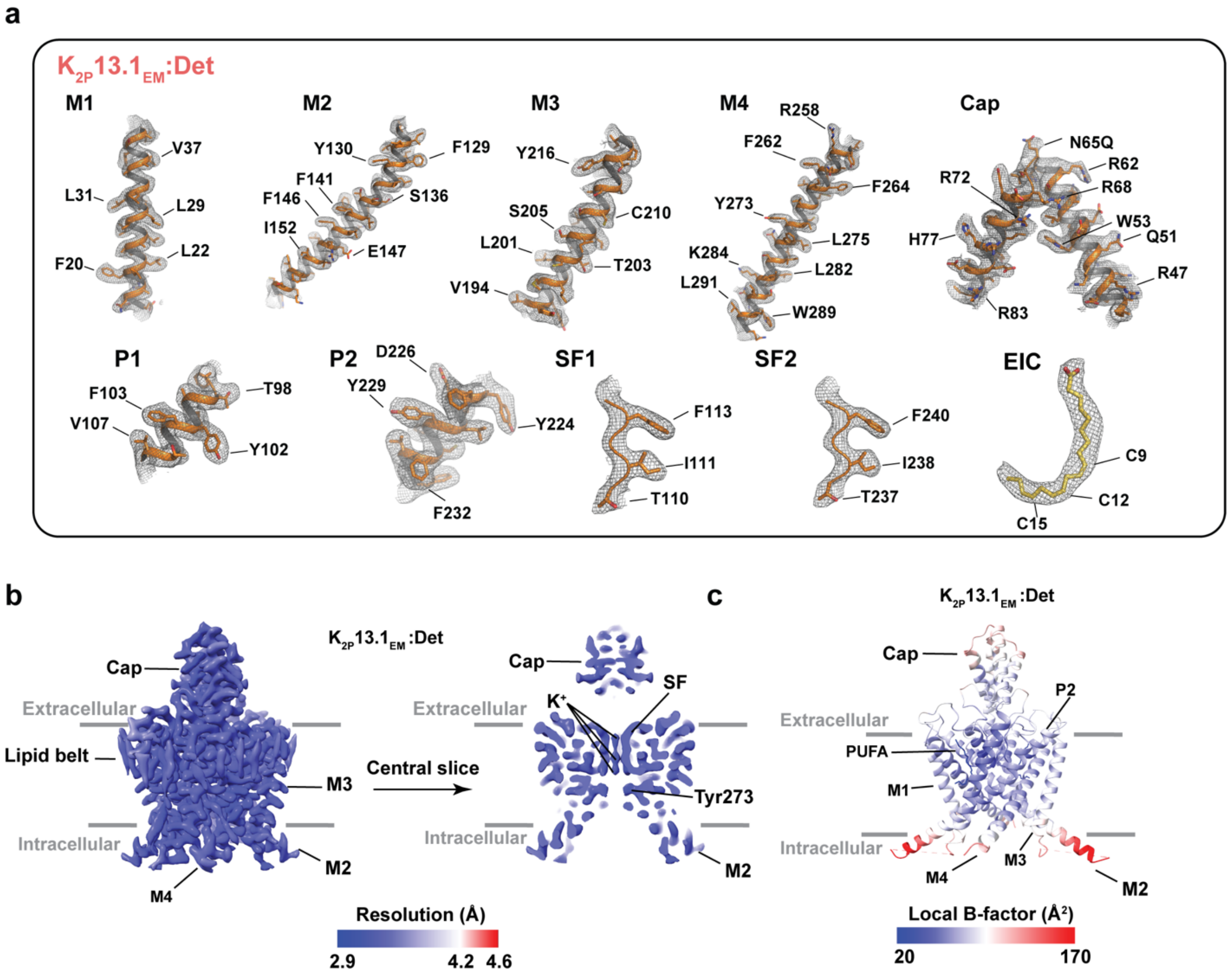


**Figure S9 $K_{2P}13.1_{EM}$:DDM:CHS:GDN (Det) complex cryo-EM map and model quality.** **a,** Cryo-EM maps for indicated $K_{2P}13.1_{EM}$ elements. Select residues are indicated. Channel elements are orange. EIC is yelloworange. Maps are rendered at 7-9σ. **b,** $K_{2P}13.1_{EM}$:Det local resolution showing a central slice through. Select channel elements and lipid belt are labeled. **c,** $K_{2P}13.1_{EM}$:Det local B-factor.

#### Figure S10 Roy-Chowdhury, *et al.*

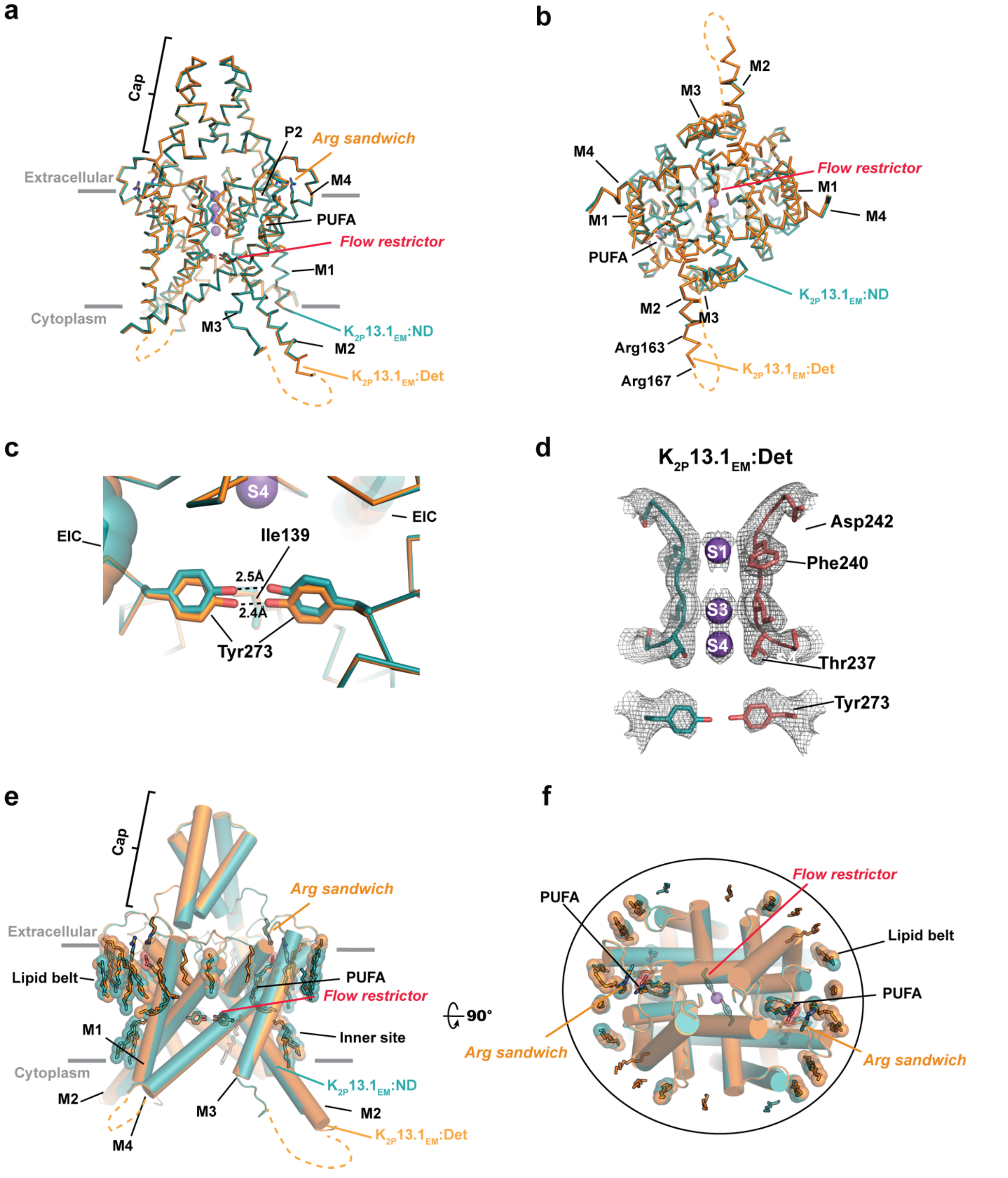


**Figure S10 Structural analysis of K2P13.1EM in nanodiscs and detergent micelles.** Superposition of K2P13.1EM:DDM:CHS:GDN, K2P13.1EM:Det (orange) and K2P13.1EM:Nanodisc, K2P13.1EM:ND (deep teal) structures showing **a,** side and **b,** cytoplasmic views. **c,** Comparison of

$K_{2P}13.1_{EM}$:Det (orange) and $K_{2P}13.1_{EM}$:ND (deep teal) flow restrictor regions. Linoleic acid (EIC) is shown in space filling. **d,** $K_{2P}13.1_{EM}$:DDM:CHS:GDN (deep teal and salmon) selectivity filter and flow restrictor densities (9$\sigma$). Select residues are labeled. Potassium ions are purple spheres. **e,** Comparison of modeled lipid positions for $K_{2P}13.1_{EM}$:Det (orange) $K_{2P}13.1_{EM}$:ND (deep teal). Common lipid positions are shown in space filling. Lipid belt, inner site lipid, and PUFA are indicated. Select channel elements are labelled. **f,** Extracellular view of ‘e’. Circle indicates lipid belt boundary. Potassium ions shown as purple spheres.

Figure S11 Roy-Chowdhury, et al.

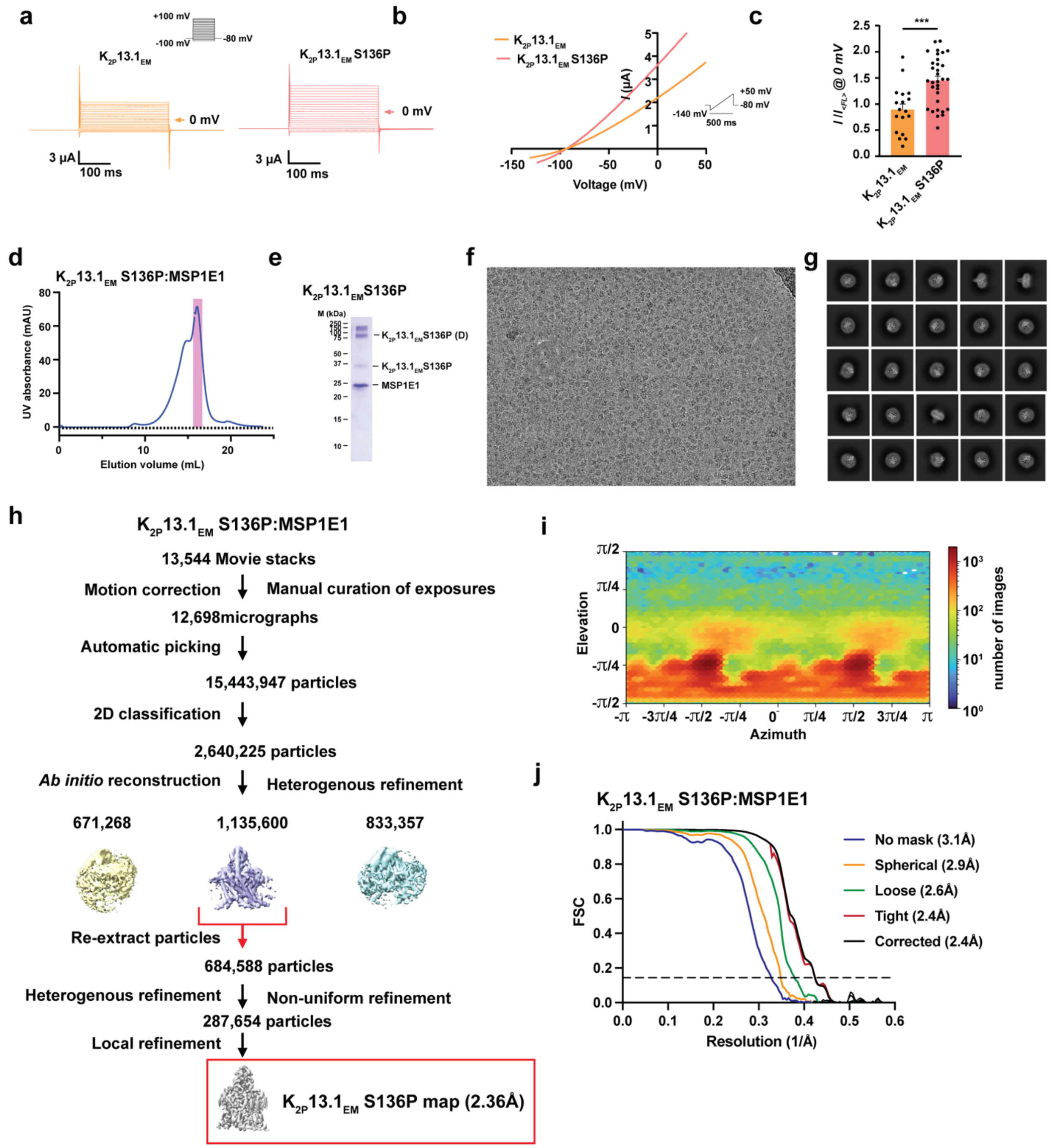


**Figure S11 Functional characterization of $K_{2P}13.1_{EM}$ S136P and cryo-EM analysis in MSP1E1 nanodiscs. a**, Exemplar TEVC recordings of $K_{2P}13.1_{EM}$ (orange) and $K_{2P}13.1_{EM}$ S136P (salmon) in *Xenopus* oocytes. 0 mV trace is indicated. Inset shows protocol. **b,** Representative TEVC current-voltage responses for $K_{2P}13.1_{EM}$ (orange) and $K_{2P}13.1_{EM}$ S136P (salmon). Inset shows protocol. **c,** Average currents $I/I_{<FL>}$ at 0 mV for $K_{2P}13.1_{EM}$ (orange) and $K_{2P}13.1_{EM}$ S136P (salmon) where $I_{<FL>}$ = average currents for full length $K_{2P}$13.1 (THIK-1). *** $p < 0.001$. Statistical analysis was performed using Kruskal-Wallis test (nonparametric ANOVA) followed by Dunn's multiple comparisons test. **d,** Exemplar SEC (Superose 6 Increase 10/300 GL) for $K_{2P}13.1_{EM}$ S136P:MSP1E1 nanodiscs. **e,** peak fraction SDS-PAGE. **f,** electromicrograph (~105,000x magnification), and **g,** 2D class averages. **h,** Workflow for cryo-EM data processing for the $K_{2P}13.1_{EM}$:MSP1E1 nanodisc complex in cryoSPARC-3.2[1]. Red arrow indicates the class of particles extracted without Fourier cropping after the initial cleanup. These were further subjected to heterogeneous, non-uniform, and local refinement jobs that resulted in the final map at 2.36Å. (red box). **i,** Particle distribution plot and **j,** gold-standard Fourier Shell Correlation (FSC) curves for the $K_{2P}13.1_{EM}$ S136P:MSP1E1 nanodisc complex.

**Figure S12** **Roy-Chowdhury, *et al.***

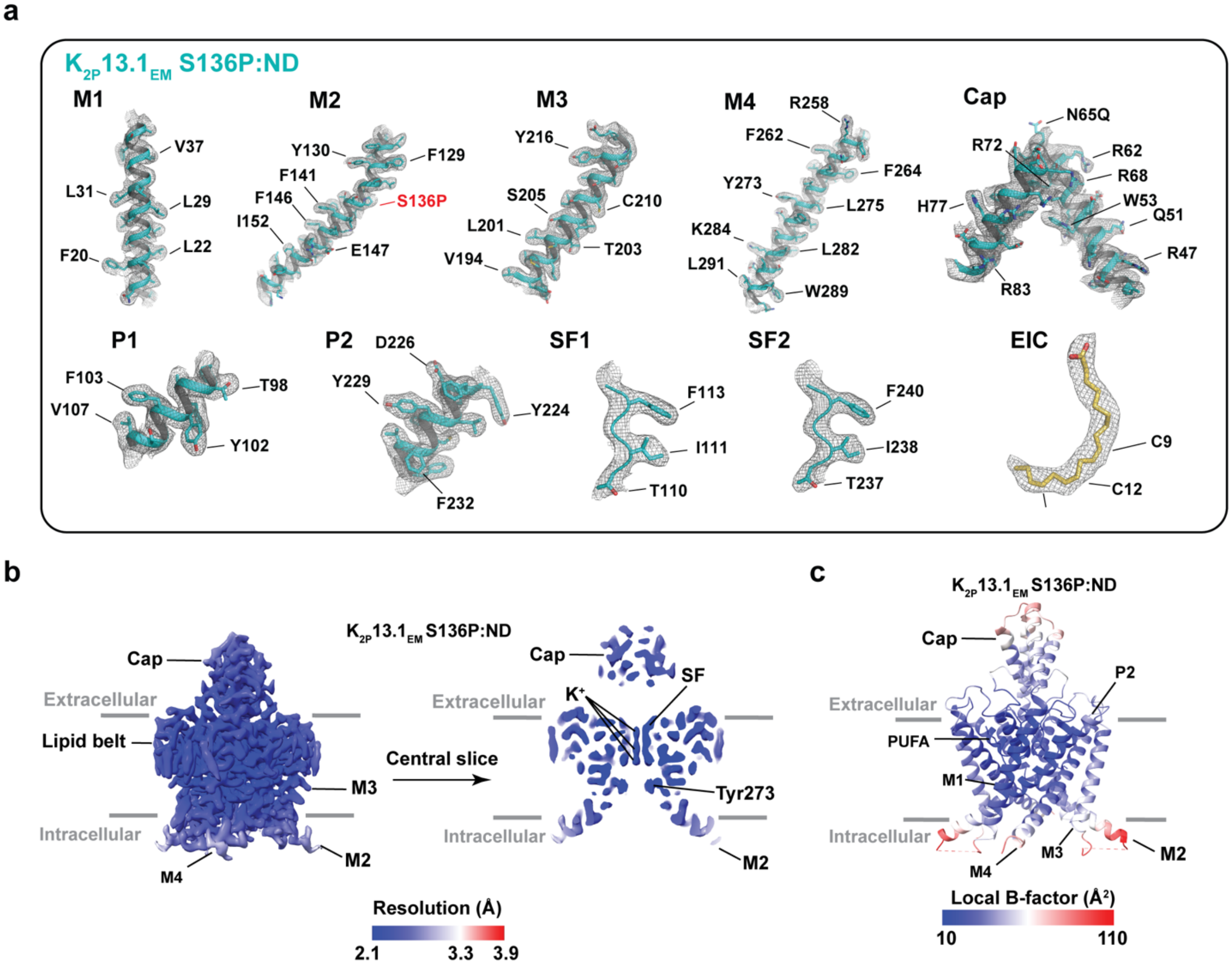


**Figure S12 K2P13.1EM S136P:MSP1E1 nanodisc (ND) complex cryo-EM map and model quality. a,** Cryo EM maps for indicated K2P13.1EM S136P elements. Select residues are indicated. Channel elements are cyan. EIC is yelloworange. Maps are rendered at 6-8σ. **b,** K2P13.1EM S136P:ND local resolution showing a central slice through. Select channel elements and lipid belt are labeled. **c,** K2P13.1EM S136P:ND local B-factor.

**Figure S13** **Roy-Chowdhury, *et al.***

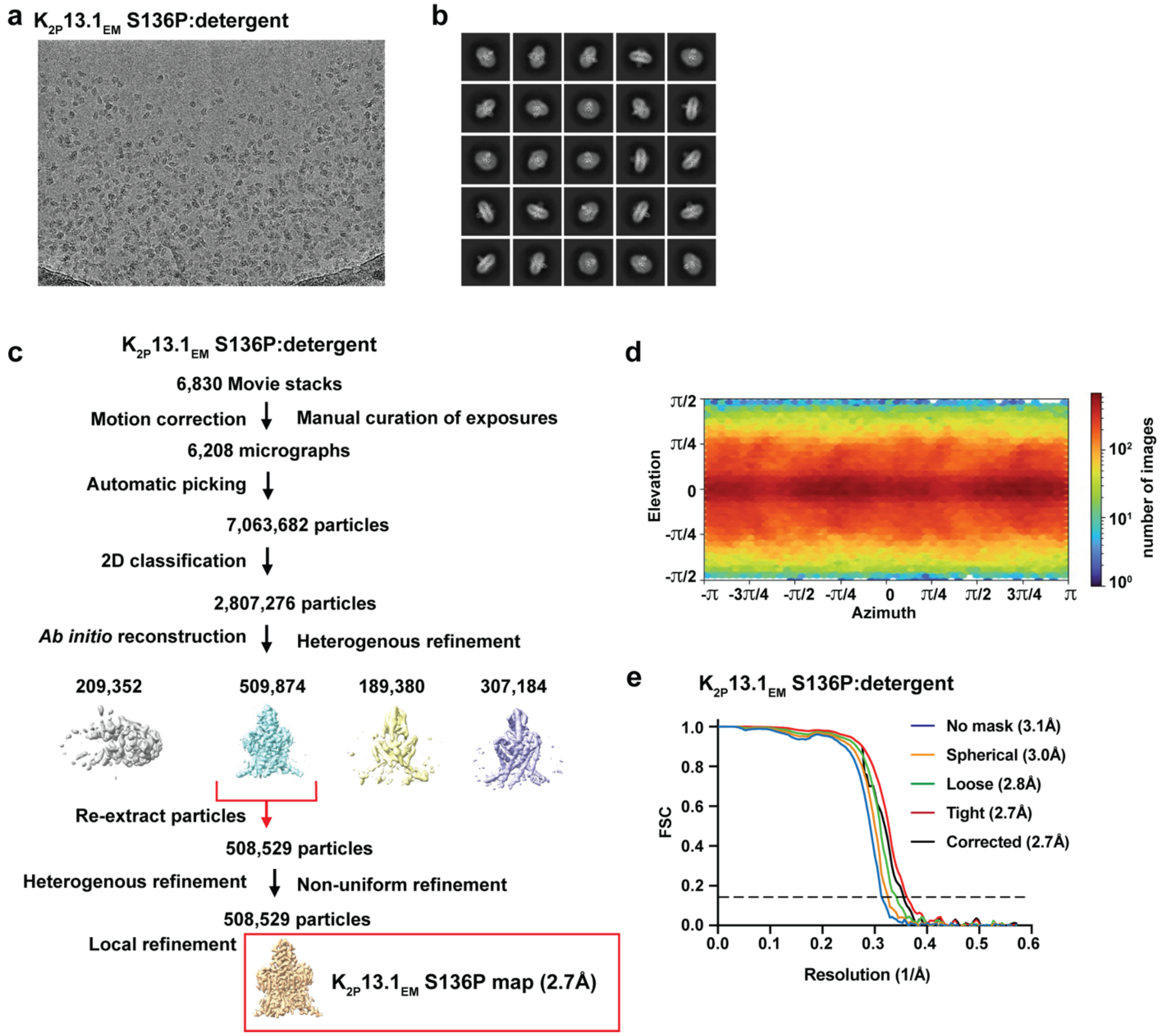


**Figure S13 Cryo-EM analysis of $K_{2P}13.1_{EM}$ S136P DDM:CHS:GDN detergent micelles**. Exemplar **a,** electromicrograph (~105,000x magnification), and **b,** 2D class averages. **c,** Workflow for cyro-EM data processing for the $K_{2P}13.1_{EM}$ S136P:DDM:CHS:GND detergent micelle complex in cryoSPARC-3.2[1]. Red arrow indicates the class of particles extracted without Fourier cropping after the initial cleanup. These were further subjected to heterogeneous, non-uniform, and local refinement jobs that resulted in the final map at 2.95Å. (red box). **d,** Particle distribution plot and **e,** gold-standard Fourier Shell Correlation (FSC) curves for the $K_{2P}13.1_{EM}$ S136P:DDM:CHS:GND detergent micelle complex.

**Figure S14** **Roy-Chowdhury, *et al.***

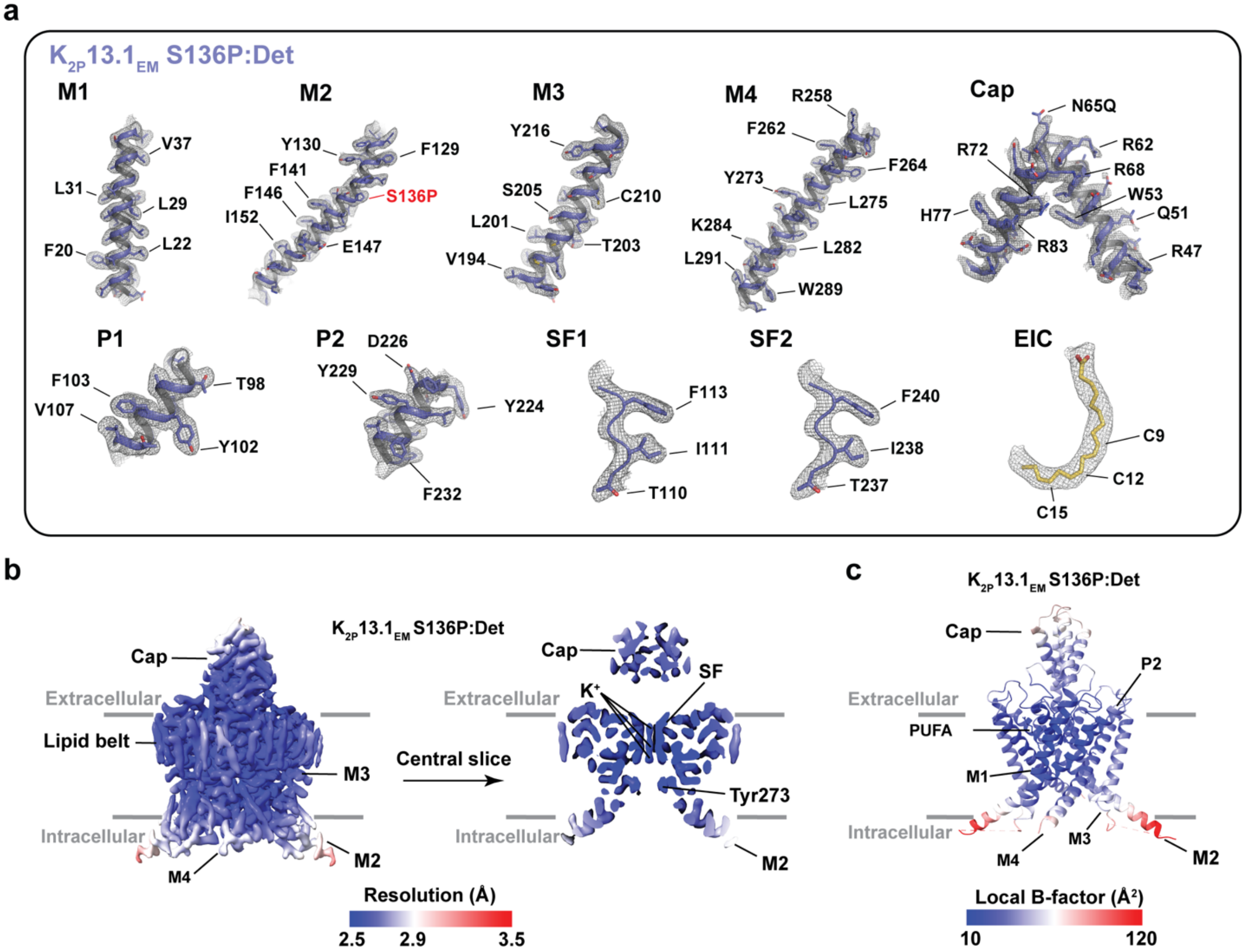


**Figure S14 $K_{2P}$13.1$_{EM}$ S136P:** DDM:CHS:GND **(Det) complex cryo-EM map and model quality. a,** Cryo-EM maps for indicated $K_{2P}$13.1$_{EM}$ S136P elements. Select residues are indicated. Channel elements are deep blue. EIC is yelloworange. Maps are rendered at 5-7σ. **b,** $K_{2P}$13.1$_{EM}$ S136P:Det local resolution showing a central slice through the channel. Select channel elements and lipid belt are labeled. **c,** $K_{2P}$13.1$_{EM}$ S136P:Det local B-factor.

**Figure S15** **Roy-Chowdhury, *et al.***

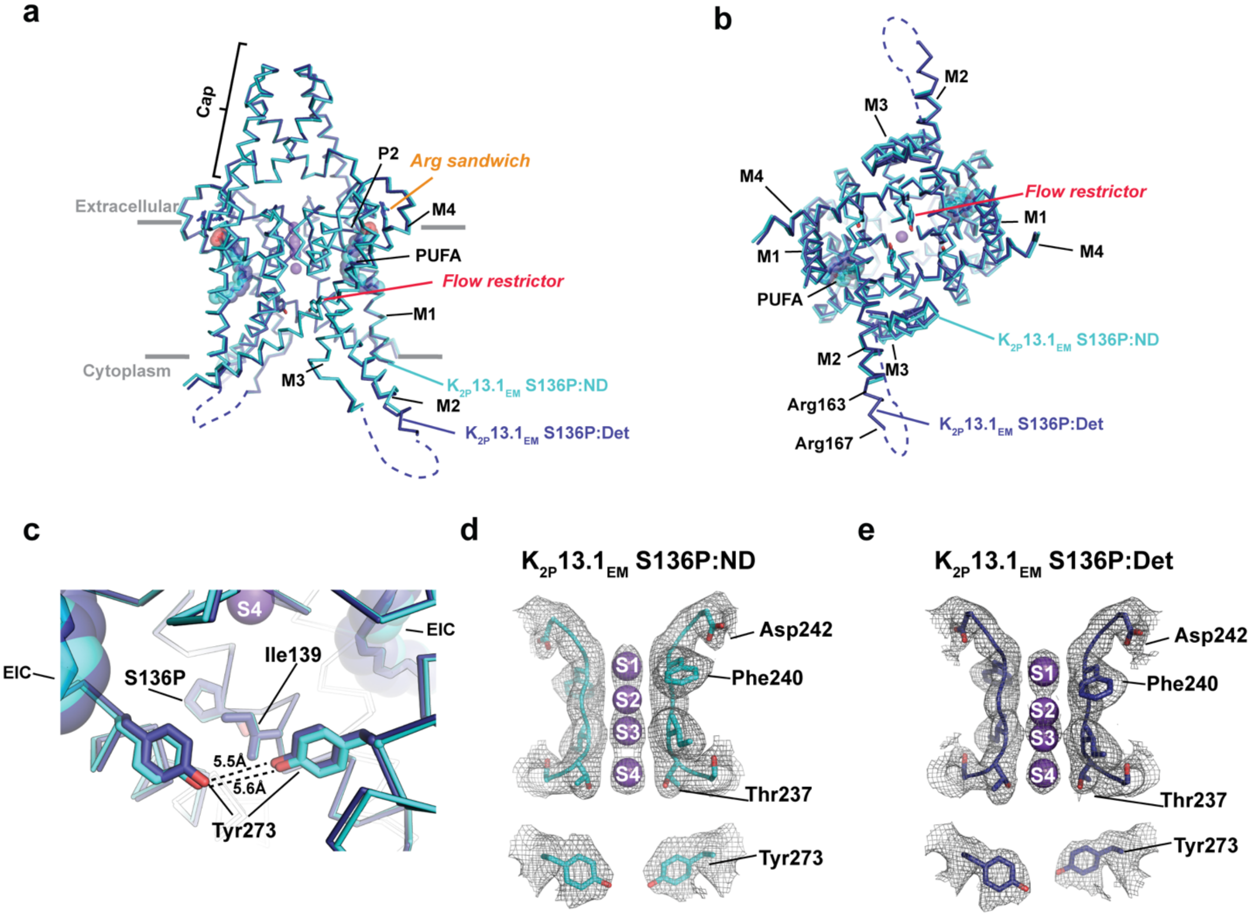


**Figure S15 Structural comparison of $K_{2P}13.1_{EM}$ in nanodiscs and detergent micelles** Superposition of $K_{2P}13.1_{EM}$ S136P:DDM:CHS:GDN, $K_{2P}13.1_{EM}$ S136P:Det (deep blue) and $K_{2P}13.1_{EM}$ S136P:Nanodisc, $K_{2P}13.1_{EM}$ S136P:ND (cyan) structures showing **a,** side and **b,** cytoplasmic views. **c,** Comparison of $K_{2P}13.1_{EM}$ S136P:ND (cyan) and $K_{2P}13.1_{EM}$ S136P:Det (deep blue) flow restrictor regions. Distances between Tyr273 hydroxyl oxygens are indicated. Linoleic acid (EIC) is shown in space filling. **d,** and **e,** Selectivity filter and flow restrictor densities (7–7.5σ) for **d,** $K_{2P}13.1_{EM}$ S136P:ND (cyan) and **e,** $K_{2P}13.1_{EM}$ S136P:Det. Select residues are labeled.

**Figure S16** **Roy-Chowdhury, *et al.***

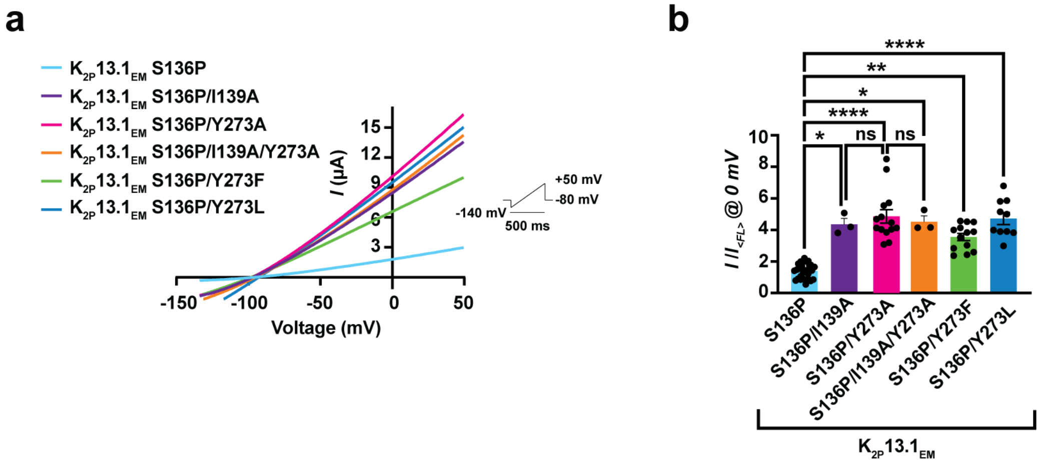


**Figure S16 Functional studies of $K_{2P}13.1_{EM}$ S136P mutants a,** Representative TEVC current-voltage responses for $K_{2P}13.1_{EM}$ S136P (cyan), $K_{2P}13.1_{EM}$ S136P/I139A (purple), $K_{2P}13.1_{EM}$ S136P/Y273A (magenta), $K_{2P}13.1_{EM}$ S136P/I136A/Y273A (orange), $K_{2P}13.1_{EM}$ S136P/Y273F (green), and $K_{2P}13.1_{EM}$ S136P/Y273L (blue). Inset shows protocol. **b,** Average currents $I/I_{<FL>}$ at 0 mV for constructs in 'a' where $I_{<FL>}$ = average currents for full length $K_{2P}13.1$ (THIK-1). Error bars are S.E.M.. **** $p < 0.0001$, *** $p < 0.001$ , ** $p < 0.01$, * $p = 0.01$-$0.05$, n.s. $p > 0.05$. Statistical analysis was performed using Kruskal-Wallis test (nonparametric ANOVA) followed by Dunn's multiple comparisons test.

**Figure S17** **Roy-Chowdhury, *et al.***

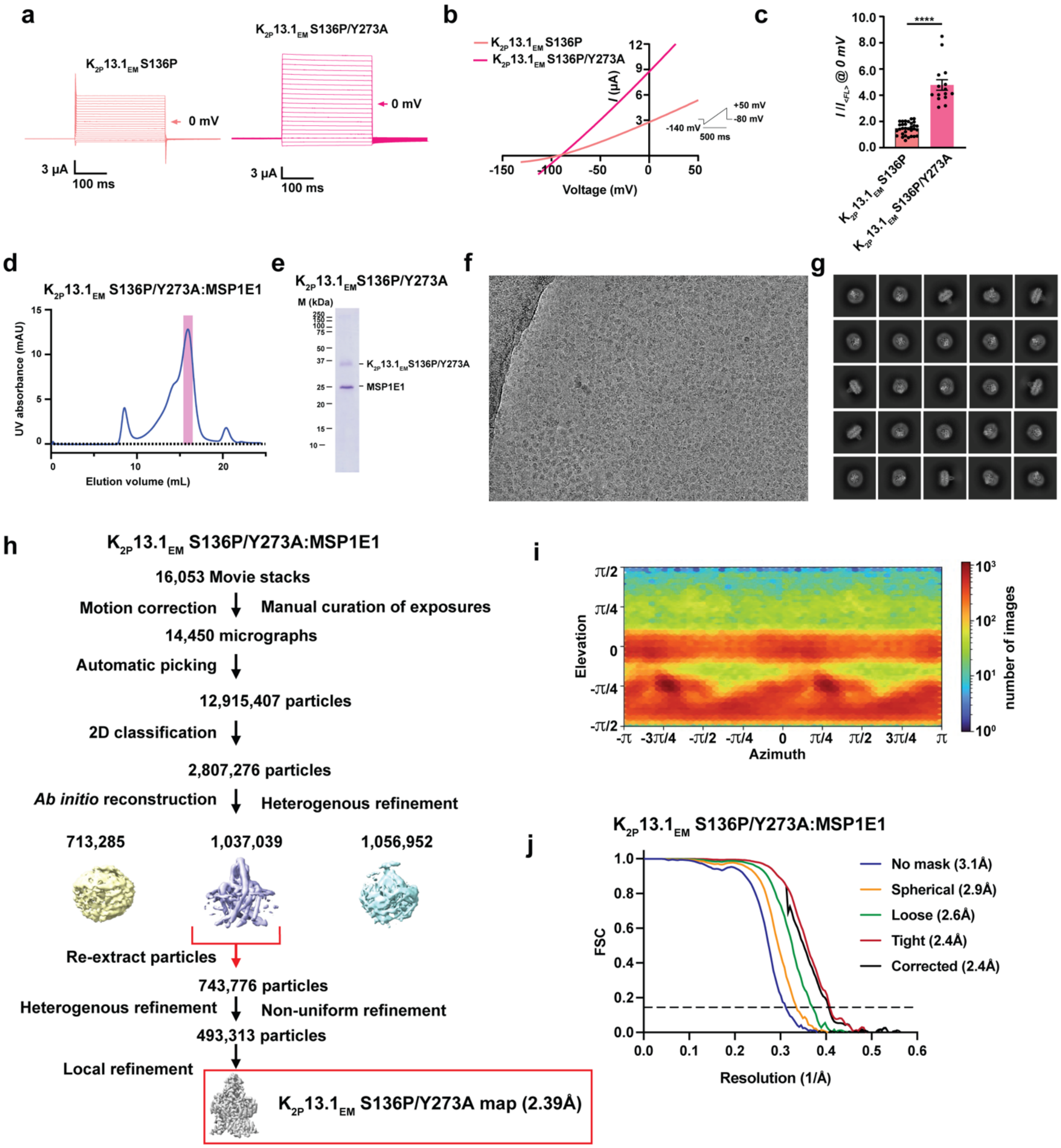


**Figure S17 Functional characterization of $K_{2P}13.1_{EM}$ S136P/Y273A and cryo-EM analysis in MSP1E1 nanodiscs. a**, Exemplar TEVC recordings of $K_{2P}13.1_{EM}$ S136P (salmon) and $K_{2P}13.1_{EM}$ S136P/Y273A (magenta) and in *Xenopus* oocytes. 0 mV trace is indicated. Inset shows protocol. **b,** Representative TEVC current-voltage responses for $K_{2P}13.1_{EM}$ S136P (salmon) and $K_{2P}13.1_{EM}$ S136P/Y273A (magenta). Inset shows protocol. **c,** Average currents $I/I_{<FL>}$ at 0 mV for $K_{2P}13.1_{EM}$

$K_{2P}13.1_{EM}$ S136P (salmon) and $K_{2P}13.1_{EM}$ S136P/Y273A (magenta) where $I_{<FL>}$ = average currents for full length $K_{2P}13.1$ (THIK-1). **d,** Exemplar SEC (Superose 6 Increase 10/300 GL) for $K_{2P}13.1_{EM}$ S136P/Y273A:MSP1E1 nanodiscs. **e,** peak fraction SDS-PAGE. **f,** electromicrograph (~105,000x magnification), and **g,** 2D class averages. **h,** Workflow for cyro-EM data processing for the $K_{2P}13.1_{EM}$:MSP1E1 nanodisc complex in cryoSPARC-3.2[1]. Red arrow indicates the class of particles extracted without Fourier cropping after the initial cleanup. These were further subjected to heterogeneous, non-uniform, and local refinement jobs that resulted in the final map at 2.39Å (red box). **i,** Particle distribution plot and **j,** Gold standard Fourier Shell Correlation (FSC) curves for the $K_{2P}13.1_{EM}$ S136P/Y273A:MSP1E1 nanodisc complex.

**Figure S18** **Roy-Chowdhury, *et al.***

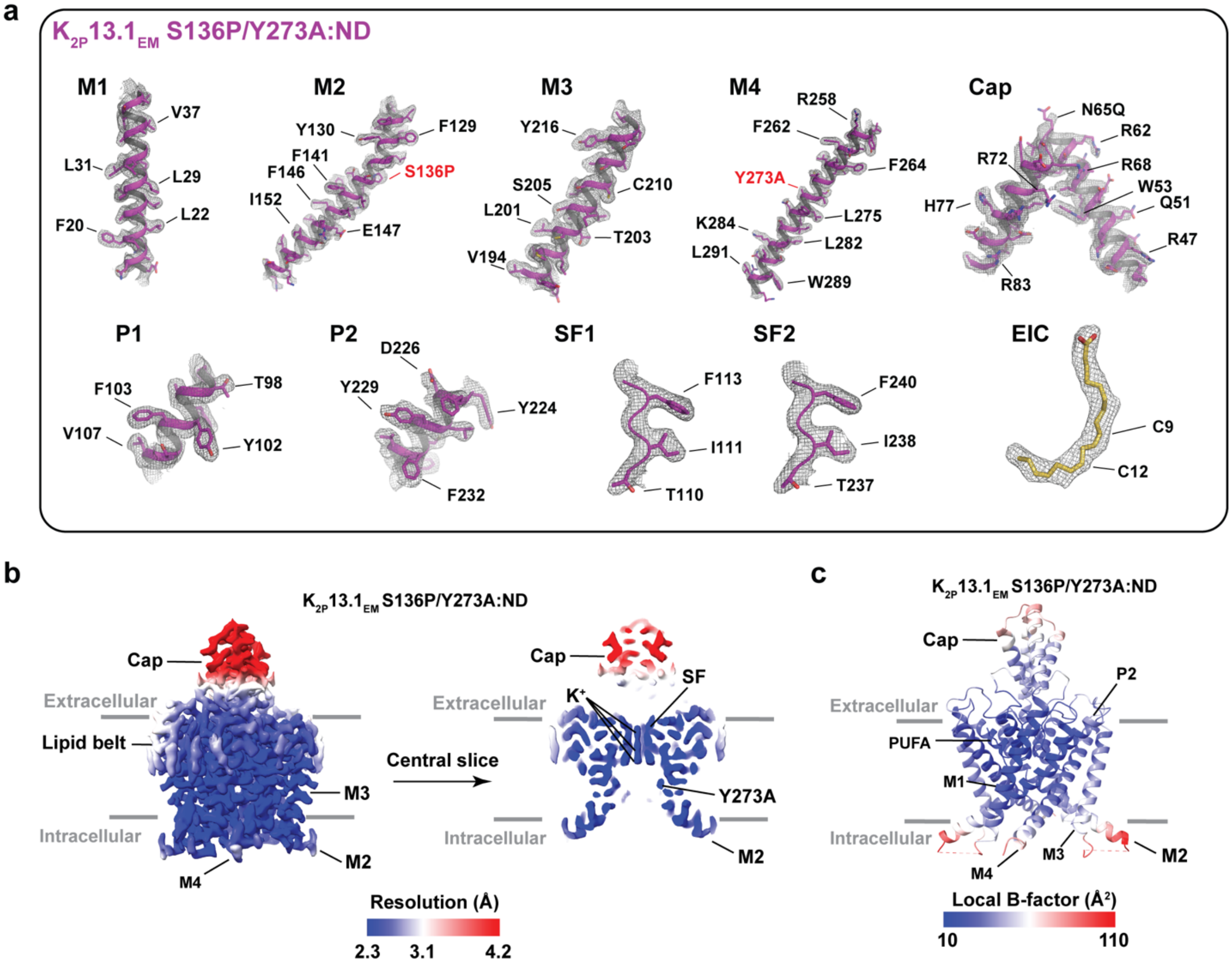


**Figure S18 $K_{2P}13.1_{EM}$ S136P/Y273A:MSP1E1 nanodisc (ND) complex cryo-EM map and model quality. a,** Cyro-EM maps for indicated $K_{2P}13.1_{EM}$ S136P/Y273A elements. Select residues are indicated. Channel elements are magenta. EIC is yelloworange. Maps are rendered at 6-8σ. **b,** $K_{2P}13.1_{EM}$ S136P/Y273A:ND local resolution showing a central slice through the channel. Select channel elements and lipid belt are labeled. **c,** $K_{2P}13.1_{EM}$ S136P/Y273A:ND local B-factor.

**Figure S19** **Roy-Chowdhury, *et al.***

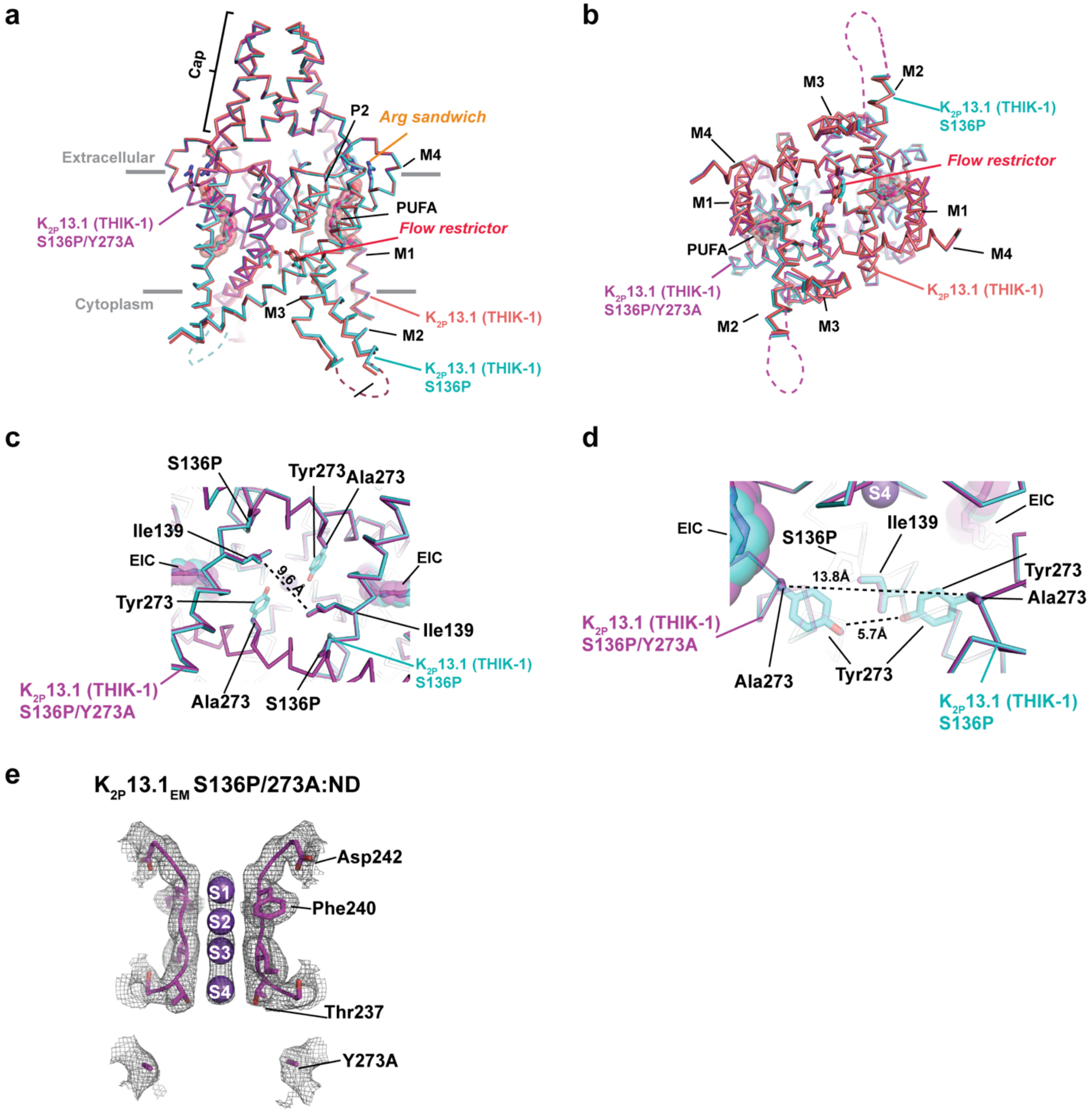


**Figure S19 K2P13.1EM S136P/Y273A:ND structural comparisons. a,** and **b,** Superposition of K2P13.1EM S136P/Y273A:ND (magenta), K2P13.1EM S136P:ND (cyan) and K2P13.1EM:ND (salmon) **a,** side and **b,** cytoplasmic views. **c,** Comparison of K2P13.1EM S136P/Y273A:ND (magenta), K2P13.1EM S136P:ND (cyan). Distance between Ile139 closest approach is indicated. **d,** Comparison of K2P13.1EM S136P:ND (cyan) and K2P13.1EM S136P/Y273A:ND (magenta) flow restrictor regions. Distances between Ala273 methyl groups and Tyr273 hydroxyls are indicated.

EIC is shown in space filling. **e,** Selectivity filter and flow restrictor densities (8σ) for $K_{2P}13.1_{EM}$ S136P/Y273A:ND (magenta). Select residues are labeled. Potassium ions are shown as purple spheres.

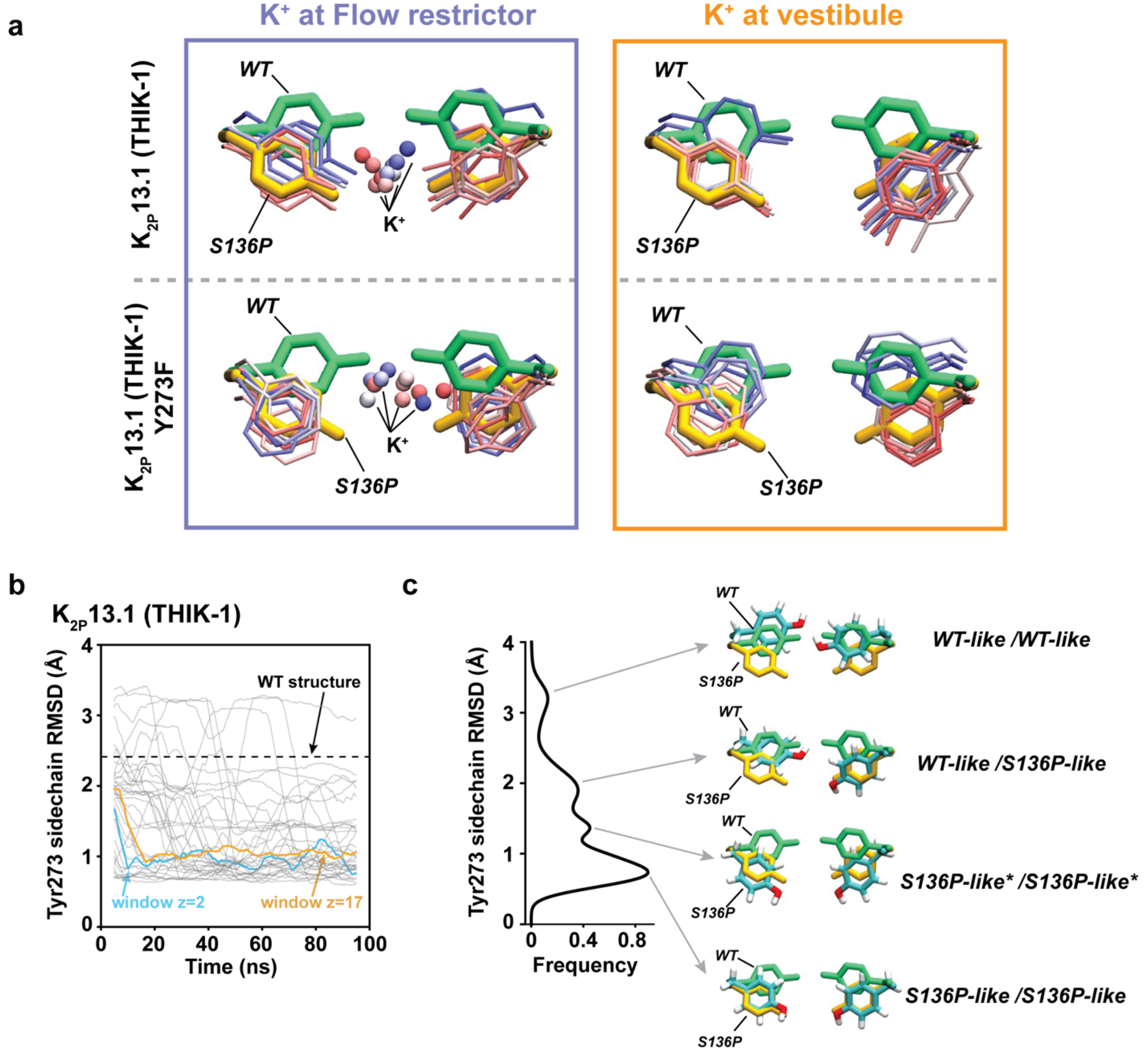


**Figure S20 Constricted flow restrictors populate the S136P state. a,** Rotamer side chain positions for Tyr273 (top) or Y273F (bottom) every 10 ns over the 100 ns umbrella sampling simulation starting from the $K_{2P}13.1_{EM}$ structure (PDB:9BSN) having the potassium ion restrained in the middle of the flow restrictor (left, window z=2) or far from the flow restrictor near the cytoplasm (right, window z=-17). The Y273 positions observed in $K_{2P}13.1_{EM}$ (PDB:9BSN) and $K_{2P}13.1_{EM}$ S136P (PDB:9C09) are shown in green and yellow, respectively. Simulation residues are colored from blue to red as time increases. **b,** Time evolution of the Tyr273 position over all umbrella sampling simulations measured by RMSD to the Tyr273 rotamer from $K_{2P}13.1_{EM}$ S136P

(PDB:9C09). Most simulations show a drop in RMSD indicating that the side chain adopts the $K_{2P}13.1_{EM}$ S136P conformation regardless of where the ion is restrained, cf. ion placed at the flow restrictor (window z=2) or far from the restrictor (window z=-17). **c,** RMSD frequency distributions from 'b'. Representative snapshots from each peak are pictured to the right. WT-like, S136P-like of Tyr273 are indicated for each subunit. S136P-like* indicates a conformation between WT-like and S136P-like. Simulated structure is shown in cyan. Y273 positions observed in $K_{2P}13.1_{EM}$ (PDB:9BSN) and $K_{2P}13.1_{EM}$ S136P (PDB:9C09) are shown in green and yellow, respectively.

**Figure S21** **Roy-Chowdhury, *et al.***

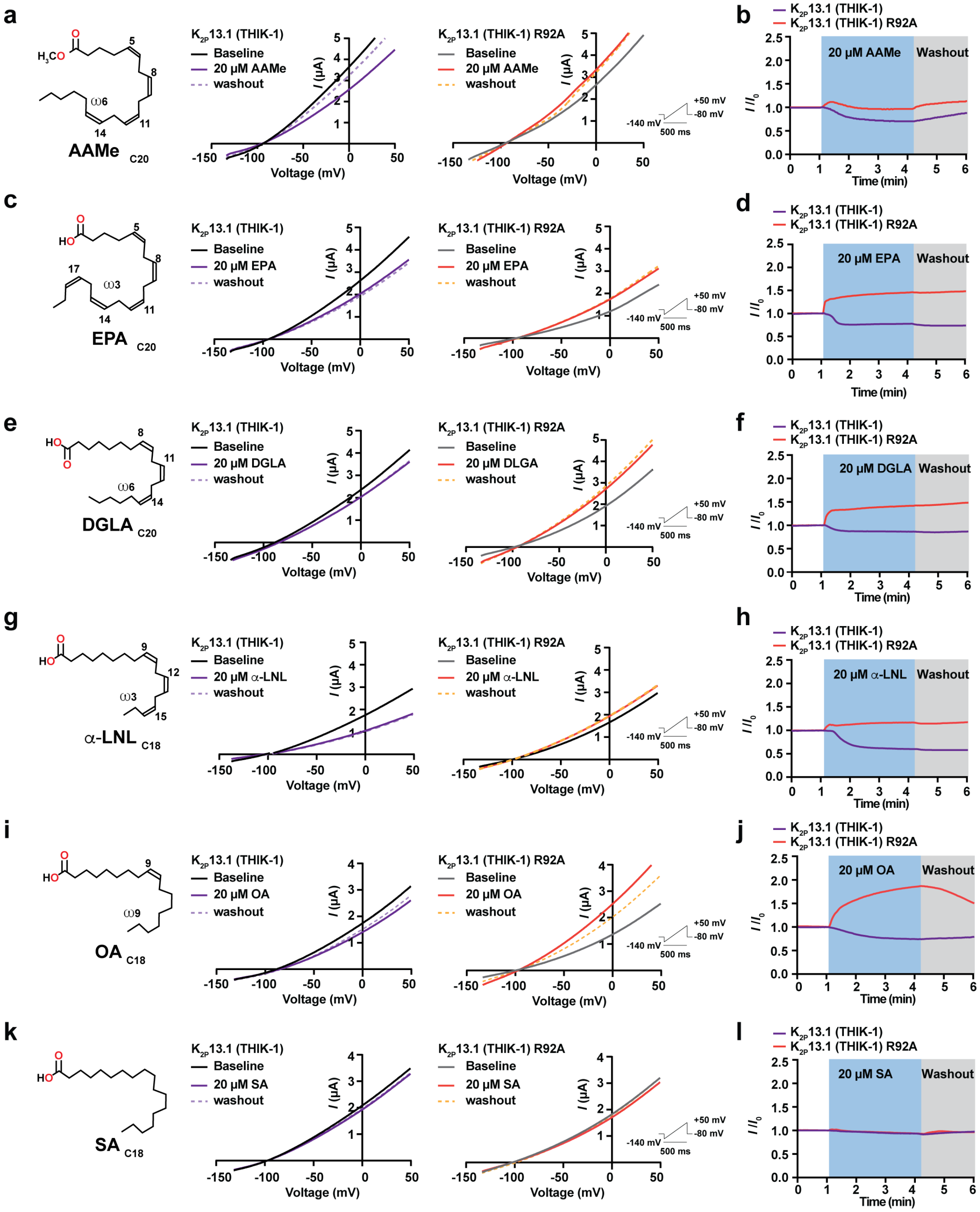


**Figure S21 K2P13.1 (THIK-1) PUFA site modulation.** Representative TEVC current-voltage responses (**a, c, e, g, i,** and **k**) and time course responses (**b, d, f**, **h, j,** and **l**) for K2P13.1 (THIK-1)

(purple) and $K_{2P}$13.1 (THIK-1) R92A (red) to **a-b,** arachidonic acid methylester (AAMe), **c-d,** eicosapentanoic acid (EPA), **e-f,** dihomo-γ-linolenic acid (DGLA), **g-h,** α-linolenic acid (α-LNL), **i-j,** oleic acid (OA), and **k-l,** stearic acid (SA). **a, c, e, g, i,** and **k** show lipid structure. Unsaturated bond locations are marked by numbers. Lipid class (ω3 or ω 6) and number of carbons are indicated. IV curves show baseline (black), lipid application (purple or red), and after 2' washout (purple or orange dashed lines).

**Figure S22** Roy-Chowdhury, *et al.*

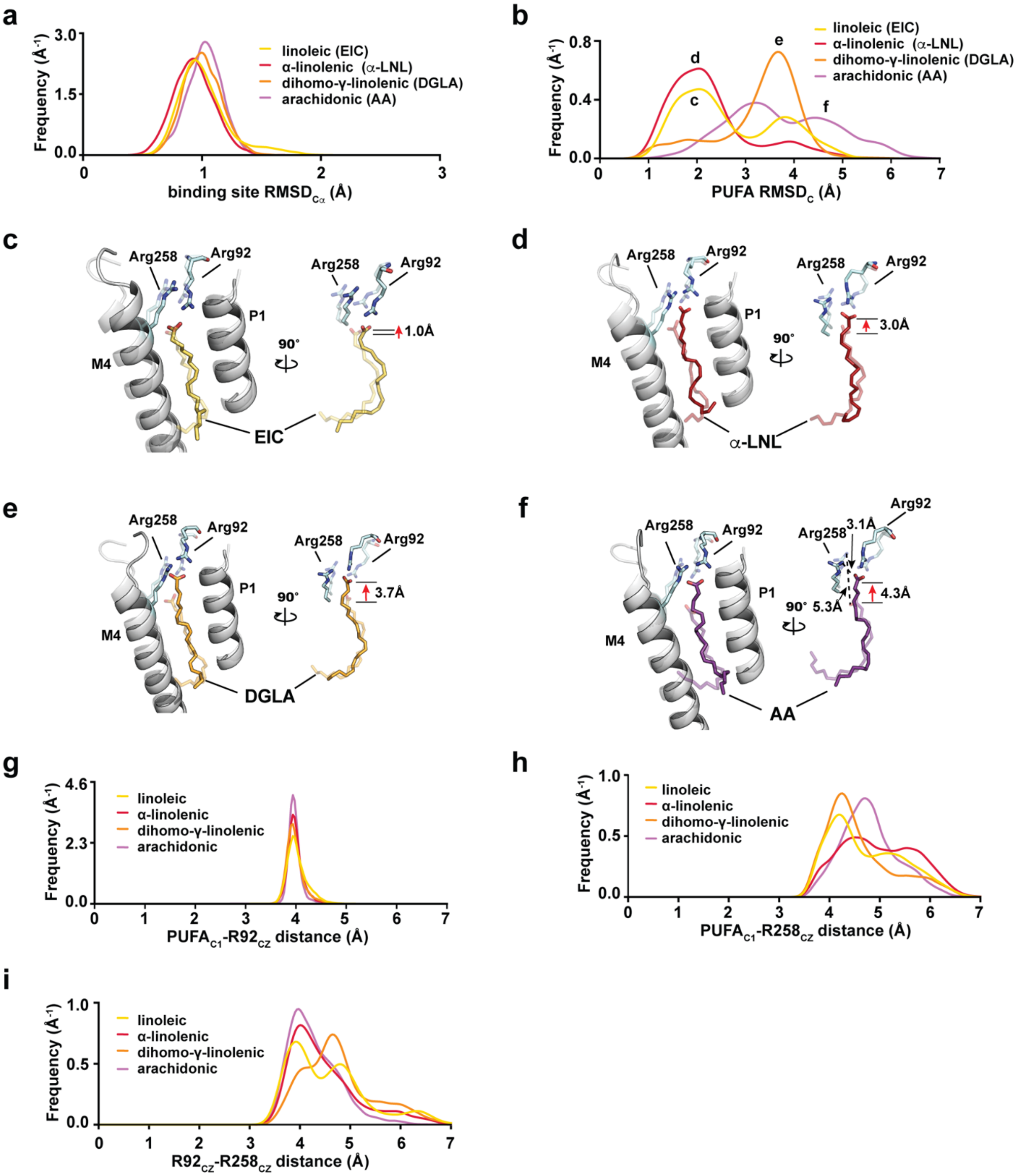


**Figure S22 $K_{2P}$13.1 (THIK-1) PUFA site simulations show binding plasticity. a,** Cα RMSD of residues lining the lipid binding pocket for each PUFA tested. **b**, PUFA RMSD with respect to the initial build (Cα residues 97-110 and 254-270). **c-f,** representative PUFA site structures from the distributions in ‘a’ for: **c,** EIC (yellow), **d,** α-LNL (firebrick), **e,** DGLA (orange), and **f,** AA (purple). In **c-f** Distances for AA carboxylate and Arg258 are indicated by the arrows. Starting structures are shown as transparent. Lipid carboxylate displacements are indicated by the red arrows. **g-i,** Observed distance distributions for: **g**, $PUFA_{C1}$-$R92_{CZ}$, **h,** $PUFA_{C1}$-$R92_{CZ}$, and **i,** $R92_{CZ}$-$R258_{CZ}$ atoms.

**Figure S23** **Roy-Chowdhury, *et al.***

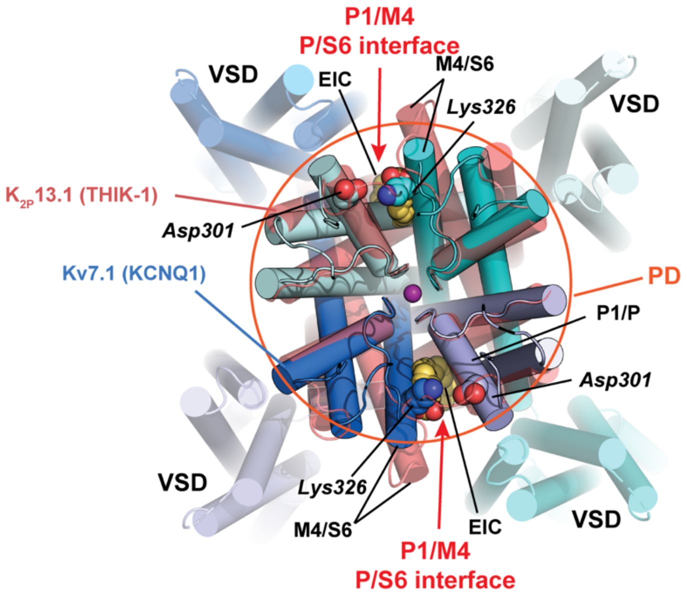


**Figure S23 $K_{2P}$13.1 (THIK-1) structural comparison with Kv7.1 (KCNQ1).** Extracellular view showing superposition of $K_{2P}$13.1 (THIK-1) (deep salmon) and Kv7.1 (KCNQ1) (marine, light blue, cyan, light cyan) (PDB:6UZZ)[9]. Lys326 and Asp301 residues framing the putative Kv7.1 (KCNQ1) PUFA site, 'Site II'[10] and EIC (yellow orange) are shown in space filling. P1/M4 and corresponding P/S6 interfaces are indicated. Circle denotes pore domain (PD). Voltage sensor domains (VSD) are labeled. Kv7.1 (KCNQ1) residues are in italics.

**Table S1 Statistics for structural data collection, refinement, and validation**

| | $K_{2P}13.1_{EM}$:ND (PDB:9BSN, EMD-44870) | $K_{2P}13.1_{EM}$:Det* (PDB:9BYI EMD-45034) | $K_{2P}13.1_{EM}$ S136P:ND (PDB:9C09, EMD-45077) | $K_{2P}13.1_{EM}$ S136P:Det (PDB:9C07, EMD-45075) | $K_{2P}13.1_{EM}$:ND S136P/Y273A (PDB:9BWS, EMD-44978) |
|---|---|---|---|---|---|
| **Data collection and processing** | | | | | |
| Magnification | 105,000 | 105,000 | 105,000 | 105,000 | 105,000 |
| Voltage (kV) | 300 | 300 | 300 | 300 | 300 |
| Electron dose (e-/Å$^2$) | 50 | 50 | 50 | 50 | 46 |
| Defocus range (μm) | -0.8 to -2.2 | -0.8 to -2.2 | -0.8 to -2.2 | -0.8 to -2.2 | -0.8 to -2.2 |
| Pixel size (Å) | 0.864 | 0.864 | 0.864 | 0.864 | 0.846 |
| Symmetry | C2 | C2 | C2 | C2 | C2 |
| Initial particle images (no.) | 11, 542, 687 | 8,081402 | 15, 443, 947 | 4,226,065 | 12, 915, 407 |
| Final particle images (no.) | 287, 654 | 375,069 | 446, 461 | 508,529 | 493, 313 |
| Map resolution (Å) | 2.65 | 2.95 | 2.36 | 2.73 | 2.39 |
| FSC threshold | 0.143 | 0.143 | 0.143 | 0.143 | 0.143 |
| Map resolution range (Å) | 2.4~3.9 | 2.95~4.6 | 2.1~3.9 | 2.0~3.8 | 2.3~4.2 |
| **Refinement** | | | | | |
| Model resolution (Å) | 2.7 | 3.2 | 2.9 | 2.8 | 2.8 |
| FSC threshold | 0.5 | 0.5 | 0.5 | 0.5 | 0.5 |
| Map sharpening *B* factor (Å$^2$) | -92.25 | -113.15 | -53.74 | -85.51 | -93.56 |
| Model composition | | | | | |
| Non-hydrogen atoms | 4346 | 4505 | 4300 | 4512 | 4286 |
| Protein residues | 514 | 524 | 514 | 524 | 514 |
| Ligands | 26 | 31 | 22 | 36 | 22 |
| *B* factors (Å$^2$) | | | | | |
| Protein | 49.00 | 80.71 | 41.29 | 43.05 | 41.28 |
| Ligand | 20.34 | 20.47 | 20.60 | 20.31 | 20.60 |
| R.m.s deviations | | | | | |
| Bond lengths (Å) | 0.004 | 0.006 | 0.004 | 0.006 | 0.005 |
| Bond angles (°) | 0.620 | 0.567 | 0.634 | 0.687 | 0.663 |
| Validation | | | | | |
| MolProbity score | 1.29 | 1.47 | 1.45 | 1.62 | 1.45 |
| Clashscore | 2.23 | 2.85 | 3.43 | 6.71 | 3.08 |
| Poor rotamers (%) | 0.92 | 0.00 | 0.69 | 0.23 | 0.93 |
| Ramachandran plot | | | | | |
| Favored (%) | 95.85 | 94.19 | 95.45 | 95.93 | 95.26 |
| Allowed (%) | 4.15 | 5.81 | 4.55 | 4.07 | 4.74 |
| Disallowed (%) | 0.00 | 0.00 | 0.00 | 0.00 | 0.00 |

* All structures were dephosphorylated forms, except $K_{2P}13.1_{EM}$:Det

**Table S2: Simulation table.** Simulation times represent the length of each window

| Channel | PUFA site lipid | Unbiased MD | Umbrella sampling |
|---|---|---|---|
| $K_{2P}$13.1 (THIK-1) | ACD | 3 x 200 ns | |
| | LAX | 3 x 200 ns | |
| | LNL | 3 x 200 ns | |
| | EIC | 3 x 200 ns | 1 x 100 ns |
| $K_{2P}$13.1 (THIK-1) Y273F | EIC | | 1 x 100ns |
| $K_{2P}$13.1 (THIK-1) Y273A | EIC | | 1 x 100ns |
| $K_{2P}$13.1 (THIK-1) S136P | EIC | | 1 x 100ns |
| $K_{2P}$13.1 (THIK-1) S136P/Y273F | EIC | | 1 x 100ns |
| $K_{2P}$13.1 (THIK-1) S136P/Y273A | EIC | | 1 x 100ns |

ACD, arachidonic acid; LAX dihomo-γ-linoleic acid; LNL α-linolenic acid; EIC, linoleic acid.

**Movie S1 $K_{2P}$13.1 (THIK-1) structural changes.** Side view of a morph between $K_{2P}$13.1 (THIK-1):ND and $K_{2P}$13.1 (THIK-1) S136P:ND structures. $K_{2P}$13.1 (THIK-1) chains are cyan and salmon. Tyr273 is shown as sticks. Potassium ions are purple spheres. Site of S136P mutation is indicated in yellow. Nanodisc is transparent.

**Movie S2 Cytoplasmic view of $K_{2P}$13.1 (THIK-1) structural changes.** Cytoplasmic view of a morph between $K_{2P}$13.1 (THIK-1):ND and $K_{2P}$13.1 (THIK-1) S136P:ND structures. $K_{2P}$13.1 (THIK-1) chains are cyan and salmon. Tyr273 is shown as sticks. Potassium ions are purple spheres. Site of S136P mutation is indicated in yellow. Nanodisc is transparent.

**Movie S3 Dynamics of $K_{2P}$13.1 (THIK-1) Tyr273 during umbrella sampling with potassium at the flow restrictor.** Evolution of Tyr273 sidechain position over 100ns umbrella sampling windows in which the ion is restrained in the flow restrictor (z = 2A) and sampled every 1 ns. showing Tyr273 positions from the $K_{2P}$13.1 (THIK-1):ND (green) and $K_{2P}$13.1 (THIK-1) S136P:ND (yellow) structures is shown. $K_{2P}$13.1 (THIK-1):ND is the initial structure for these simulations. In the equilibration run (not shown) performed before the 100ns production run, Tyr273 already begins to move away from the starting pose.

**Movie S4 Dynamics of $K_{2P}$13.1 (THIK-1) Y273F during umbrella sampling with potassium at the flow restrictor.** Evolution of Y273F sidechain position over 100ns umbrella sampling windows in which the ion is restrained in the flow restrictor (z = 2A) and sampled every 1 ns. showing Tyr273 positions from the $K_{2P}$13.1 (THIK-1):ND (green) and $K_{2P}$13.1 (THIK-1) S136P:ND (yellow) structures is shown. $K_{2P}$13.1 (THIK-1):ND is the initial structure for these simulations. In the equilibration run (not shown) performed before the 100ns production run, Y273F already begins to move away from the starting pose.
